# Structural Co-optation and Loss-of-function Underlie the Evolution of Regulatory Novelty in the Glucokinase Regulatory Protein

**DOI:** 10.64898/2026.05.08.723886

**Authors:** Joshua I. Santiago, Carolin Freye, S. Shirin Kamalaldinezabadi, Johanna E. Papa, A. Carl Whittington, Brian G. Miller

**Author notes:** Corresponding authors: A. Carl Whittington and Brian G. Miller, **Email:** and. Department of Chemistry and Fermentation Sciences, Appalachian State University; Boone, NC, USA. **Author Contributions:** J.I.S., A.C.W, and B.G.M designed research; J.I.S., C.F., and A.C.W. performed research; S.S.K. and J.E.P. provided materials; J.I.S., A.C.W., and B.G.M analyzed data; and J.I.S. and A.C.W. wrote the paper. **Competing Interest Statement:** The authors declare no competing interests.

## Abstract

The glucokinase regulatory protein (GKRP) derives from an ancestral etherase. Despite existing as a single locus in the metazoans, GKRP evolved multiple novel functions unrelated to etherase activity. In jawed vertebrates, a protein-protein interaction (PPI) emerged that inhibits glucokinase (GCK) activity in the liver. This PPI is critical to maintaining glucose homeostasis. In mammals, GKRP is allosterically regulated by carbohydrates, with 6-phospharylated sugars promoting inhibition of GCK by GKRP, while 1-phosphorylated sugars relieve inhibition. Here, we use a vertical evolutionary approach to identify the genetic, biochemical, and biophysical mechanisms underlying the emergence of small-molecule allostery in GKRP. We pinpointed a single leucine to valine substitution in the N-terminus of GKRP from the ancestor of the euarchontoglires that, when introduced into the non-regulated placental mammal GKRP ancestor, installed sensitivity to sorbitol-6-phosphate (S6P). Interestingly, GKRP’s inhibitory activity in the absence of S6P was reduced but unchanged in its presence. The mutation enabled co-optation of the ancestral etherase active site, which also existed as an ambiguous phosphorylated carbohydrate binding site in unregulated GKRPs. This substitution likely introduced an alternative conformation of the N-terminus causing apo-GKRP to sample a binding incompetent state prior to GCK binding. Our results suggest a simple model of the evolution of protein functional novelty where a single mutation can cause a large functional shift via co-optation of pre-existing structural features. Importantly, in contrast to many models of protein evolution, ours does not require the addition of new genetic material to realize a novel function such as small-molecule allosteric regulation.

## Introduction

The evolution of functional novelty in proteins is essential to organismal diversification (1, 2). Metabolic expansion (3, 4), cell signaling (5, 6), and protein regulation (7, 8) all require the emergence of new protein functions. Current models of the evolution of novel protein functions require the introduction of new genetic material via gene duplication (9–11), horizontal gene transfer (12, 13), or *de novo* gene birth (14, 15). At the other end of that process lies gene death, in which a previously protein-encoding gene loses its selected function and drifts away, becoming a pseudogene (16).

In contrast to these models of functional novelty, the glucokinase regulatory protein (GKRP) appears to have existed as a single locus across eukaryotic evolution (17). Despite the lack of new genetic material, GKRP has evolved at least two novel functions in the vertebrate lineage (18). The first trait to emerge was an inhibitory protein-protein interaction with glucokinase (GCK). In the liver, the synthesis of glucose-6-phosphate by GCK triggers glycogen synthesis (19). When systemic glucose levels are low, GKRP binds to GCK inhibiting its activity (20, 21). Additionally, the GCK-GKRP complex is translocated to the nucleus, removing GCK activity from the cytoplasm (22). Recently, Kamalaldinezabadi *et al*. demonstrated that this regulatory, heteromeric interaction between GKRP and GCK evolved in the ancestor of the gnathostomes (jawed vertebrates) via exaptation of a hydrophobic patch in GCK and the insertion of a binding loop into GKRP that fits into the GCK patch (18).

The second new function to emerge was allosteric regulation of GKRP’s inhibitory activity by phosphorylated carbohydrates, a characteristic that is observed in extant mammals. In the unliganded state (23), an N-terminal extension in GKRP adopts a folded conformation distal to the GCK binding site at the junction of its constituent lid and dual sugar-isomerase domains (SIS; Fig. 1A). In the bound state (20), the N-terminus an extended conformation where it can interact with residues at the GCK binding interface. In rats and humans, phosphorylated carbohydrates such as fructose-6-phosphate (F6P) and sorbitol-6-phosphate (S6P) promote the GKRP-GCK interaction and enhance inhibition. Conversely, fructose-1-phosphate (F1P) inhibits the GKRP-GCK interaction. Intriguingly, the GKRP from the frog *Xenopus laevis* is insensitive to allosteric regulation by phosphorylated carbohydrates despite the structure of the frog GKRP-GCK complex (24) being superimposable with the mammalian structure, including the positioning of bound ligands.

**Figure 1.**
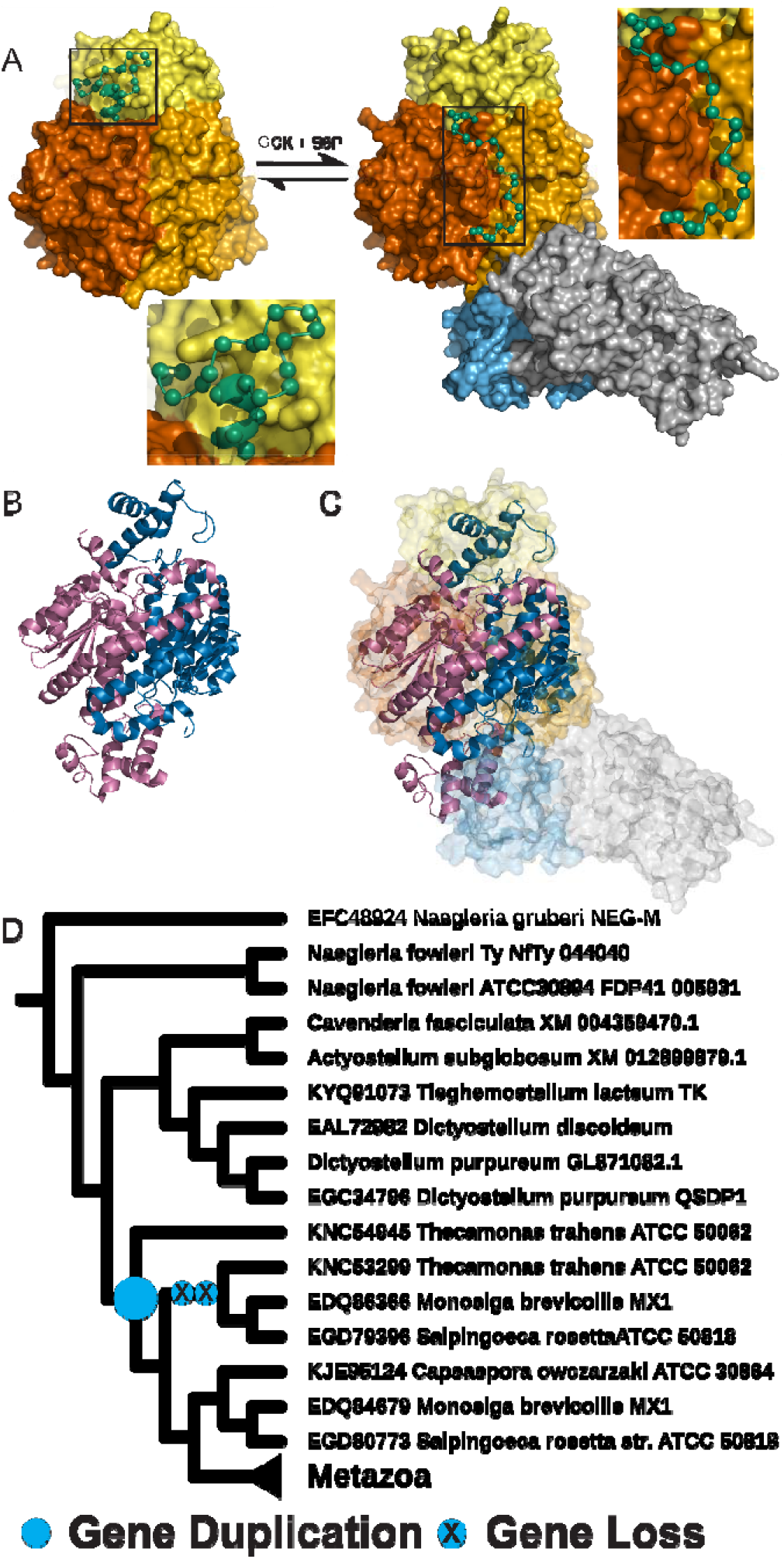
GKRP structure, function, and evolution. (*A*) GKRP (left; pdb:4BB9) consists of two SIS domains (shades of orange) and a LID domain (yellow). GKRP (right; pdb:4LC9) binds to GCK in a cleft between its large (gray) and small (light blue) domains inhibiting GCK’s ability to phosphorylate glucose. Binding of GCK is associated with a rearrangement of the GKRP N-terminus (residues 1-30; green spheres). In mammals, GKRP inhibition of GCK is allosterically modulated by phosphorylated carbohydrates with sorbitol-6-phosphate (S6P) promoting GKRP-GCK complex formation. *Insets* show a zoomed view of N-terminus secondary structure. (*B*) MurQ (pdb:4LZJ) is a bacterial etherase that hydrolyses MurNAc-6P during cell wall recycling. It functions as a homodimer (dark blue and purple). (*C*) GKRP evolved from a duplication and unequal fusion of two copies of MurQ. (*D*) Analysis of GKRP synteny among eukaryotes demonstrates that after a gene duplication event in the Apusomonad ancestor, one copy was lost prior to the divergence of Metazoa, among which GKRP generally exists as a single locus (see SI).

Eukaryotic GKRP evolved from a duplication and uneven fusion of a bacterial gene, *murq* (17), that encodes a homodimeric bacterial etherase involved in cell wall recycling (Fig. 1B) (25, 26). The structure of the MurQ homodimer (27) is superimposable with mammalian GKRP, including the active site where a MurQ inhibitor overlaps with S6P in the mammalian structure. The GKRP homolog from the excavate amoeba *Naegleria gruberi* is closer in sequence to vertebrate GKRPs than to MurQ but displays etherase activity comparable to bacterial homologs (17). GKRP has no other known role in *N. gruberi*, suggesting that GKRP plays some potential role related to MurQ-like activity in the amoeba and related organisms, likely digestion of bacterial cell walls (17).

Strangely, frog GKRP also displays etherase activity, albeit diminished by multiple orders of magnitude, despite no known role for MurQ-like activity in vertebrates.

We used phylogenetics and genomic analysis to determine orthology and synteny in eukaryotic GKRPs. We measured etherase activity and GCK inhibition in extant GKRPs as well as extinct, ancestral GKRPs that we resurrected in the lab. Combined with structural and mutational analysis, we uncovered the genetic, biochemical, and biophysical mechanisms underlying the evolution of functional novelty in GKRP as it transitioned from an active etherase to a member of a protein-protein inhibitory complex that is allosterically regulated by small molecules. Our results demonstrate a model for the evolution of functional novelty that does not require the addition of new genetic material but instead relies on co-optation and repurposing of protein features encoded at an already existing locus.

## Results and Discussion

### GKRP is a single locus among Metazoa

The functional transition of GKRP to a regulatory protein occurred during vertebrate evolution, specifically between the last common ancestor of vertebrates and the ancestor of the jawed vertebrates (18). The phylogeny of Kamalaldinezabadi *et al*. shows GKRP as a single locus across all eukaryote evolution. This suggests that no gene duplication event occurred during the evolution of eukaryotic GKRP, which could have provided the genetic raw material to facilitate the functional novelty found in GKRP in the jawed vertebrates and the small-molecule allostery found in mammals. To determine if this was a potential sampling artifact from the previously estimated GKRP phylogeny (18), we investigated GKRP genomic loci among eukaryotes.

Ensembl (28) and Genomicus (29) databases enable analysis of relationships of orthology, paralogy, and synteny through gene structure alignments and gene order alignments, respectively, integrated with gene trees. Among chordates and tunicates, Ensembl shows a single GKRP ortholog with highly conserved exon-intron architecture (Fig. S1). In the jawed vertebrates, Genomicus output indicates these are syntenic orthologs (Fig. S2 & S3). The Genomicus output demonstrates the orthology of GKRPs in vertebrates, tunicates, and cephalochordates (Fig. S2 and S3), as well as linking these orthologs to the invertebrate metazoans (Fig. S4). Among the invertebrates, there are lineage specific events, i.e. duplications in a species of sea urchin and a species of ribbon worm, and a gene split event in an anemone (Fig. S5). However, Metazoan GKRPs are generally single locus orthologs.

Among Protista, there was at least one gene duplication event prior to the divergence of Metazoa in the ancestor of Apusomonididae and Holozoa (Fig. S6). We constructed a neighbor-joining GKRP phylogeny with amino acid sequences to estimate the relatedness of metazoan GKRPs to the protist paralogs. Our GKRP tree shows that the Metazoan GKRPs cluster with only one of these early GKRP paralogs (Fig. 1D). Independent *gkrp* losses in *Capsaspora* and the ancestor of Metazoa supports our contention that GKRP has existed as a single ortholog since the divergence of animals, long before the novel functions emerged in the jawed vertebrates (Fig. 2A) (18).

**Figure 2.**
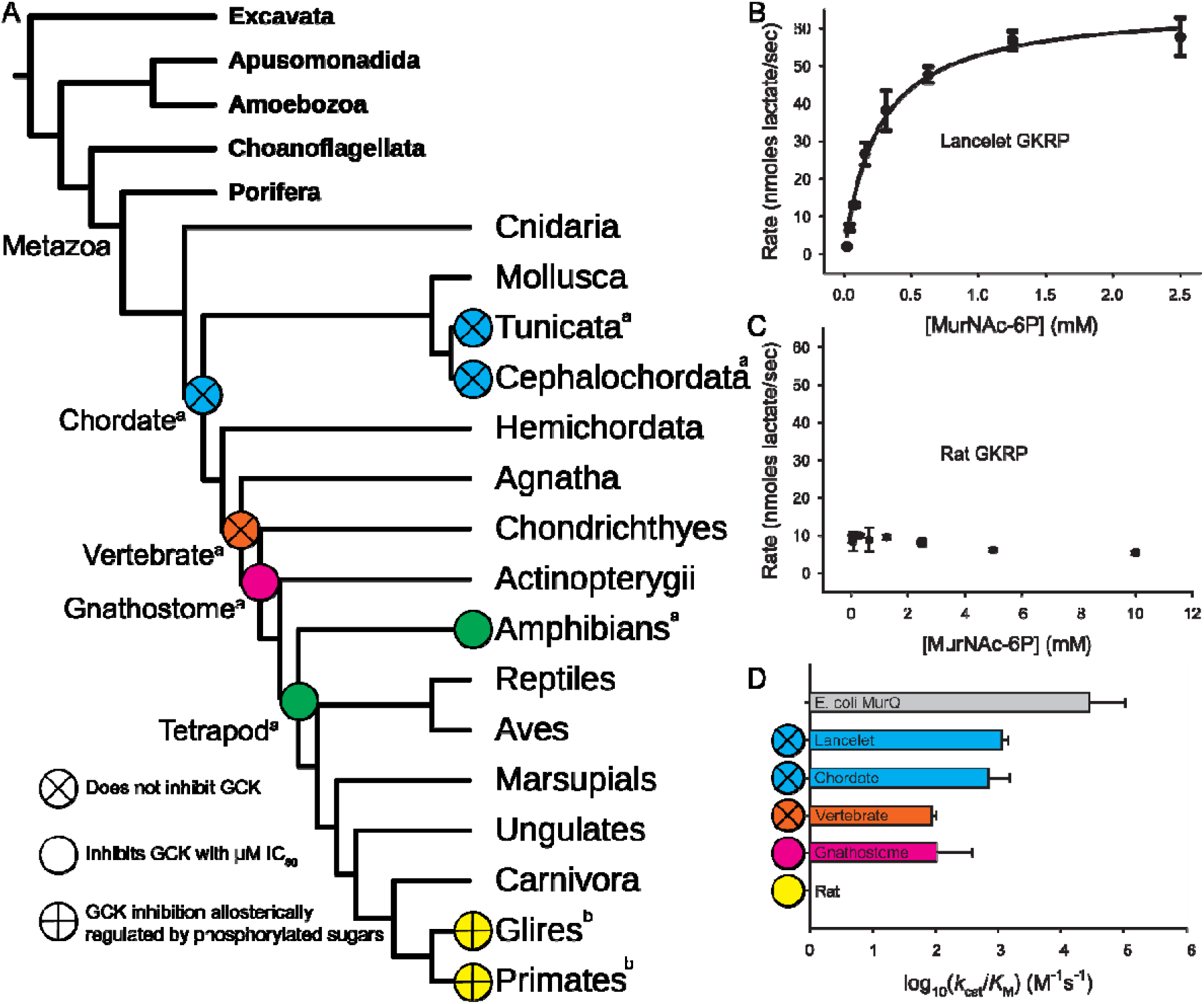
The emergence of GKRP inhibition of GCK in vertebrates and allosteric regulation of GKRP activity in mammals correlates with a loss of ancestral etherase activity. (*A*) Eukaryotic GKRP phylogeny displaying functional and regulatory properties of extant and extinct GKRPs (phylogeny adapted from (18); ^a^ – data from (18); ^b^ – data from (30). (*B*) Representative etherase activity curve showing Michaelis-Menten kinetics for Lancelet GKRP. Data points represent the average observed rates from three technical replicates. Error bars represent standard deviation. (*C*) Representative etherase activity measurements for Rat GKRP showing no ability to hydrolyze MurNAc-6P. Data points represent the average observed rates from three technical replicates. Error bars represent standard deviation. (*D*) Specificity constants for MurNAc-6P for extant and extinct MurQ and GKRPs show the loss of etherase activity across GKRPs evolution. Bars represent the average of two biological replicates. Error bars represent standard deviation.

### Non-mammalian GKRPs are active etherases

The inhibitory interaction between GKRP and GCK is specific to gnathostomes, the jawed vertebrates (Fig. 2A). Extant cephalochordate and tunicate GKRP orthologs do not inhibit their corresponding GCKs (18). This correlates with the phenotype seen in the chordate and vertebrate ancestral GKRPs, which also do not inhibit their corresponding GCKs. As the ancestral function of GKRPs is etherase activity, we tested if etherase activity was compatible with the evolution of the GKRP-GCK interaction along the vertebrate trajectory. We measured etherase activity in both extant GKRPs, and the extinct ancestral GKRPs from Kamalaldinezabadi *et al*.using a linked enzyme assay coupling hydrolysis of the *N*-acetylmuramic acid 6-phosphate (MurNAc-6P) lactyl ether bond to production of D-lactate, which is coupled to formazan production. As a control and to provide a benchmark for potential etherase activity, we measured the activity of the extant MurQ from *Escherichia coli* and found that it displays strong etherase activity and is subject to substrate inhibition (3140 ± 531 M^-1^ s^-1^; *K*_i_ = 3.36 ± 0.75 mM; Fig. S7; Table S1).

A representative activity assay of the extant GKRP from the cephalochordate lancelet demonstrates that etherase activity is retained in non-vertebrate GKRPs, albeit without substrate inhibition (1493 ± 166 M^-1^ s^-1^; Fig. 2B; Fig. S8; Table S1). In contrast, rat GKRP lacks detectable etherase activity (Fig. 2C; Fig. S9; Table S1). Assays of extinct GKRPs demonstrate that the chordate (703 ± 66 M^-1^ s^-1^; Fig. 2D; Fig. S10; Table S1) and vertebrate (86 ± 6 M^-1^ s^-1^; Fig. 2D; Fig. S11; Table S1) ancestral GKRPs retained etherase activity. Further, the gnathostome ancestral GKRP also displays relatively strong etherase activity (105 ± 13 M^-1^ s^-1^; Fig. 2D; Fig. S12; Table S1) demonstrating that enzymatic activity and formation of the GKRP-GCK interaction are not mutually exclusive. This comports with the previous report of etherase activity in GKRP from the frog *Xenopus laevis* (17).

In addition to displaying etherase activity, Veiga-da-Cunha *et al*. demonstrated that the etherase activity of frog GKRP is inhibited by F6P and most strongly by S6P (17). In contrast, F1P does not inhibit the etherase activity. Those authors were able to measure small amounts of etherase activity in rat GKRP and found that it is similarly inhibited by F6P and S6P. Interestingly, it is also strongly inhibited by F1P. As stated above, structural data indicates that phosphorylated carbohydrates bind at the etherase active site (20, 24). These observations support a model in which the GKRP etherase active site likely existed as an ambiguous phosphorylated carbohydrate binding site before the allosteric modulation of GKRP inhibitory activity emerged.

### Mechanistic evolution of small-molecule allosteric regulation in GKRP

To determine when along the vertebrate evolutionary trajectory allosteric activation of GKRP inhibitory activity (IC_50_) by phosphorylated carbohydrates first emerged, we measured inhibition of corresponding GCKs by extant and resurrected ancestral GKRPs in the presence and absence of S6P (Table S2). The GKRP from lancelet was previously shown to not inhibit lancelet GCK in the absence of S6P (18). Addition of S6P does not induce inhibitory activity in the lancelet pair (Fig. S13). Frog GKRP inhibits frog GCK with a sub-micromolar IC_50_ (Fig. S14) and S6P has no effect on the inhibitory activity (Fig. S15). Our resurrected ancestral GKRPs show a similar pattern. The chordate and vertebrate ancestral GKRPs do not inhibit chordate and vertebrate GCKs (18), respectively, and addition of S6P causes no change (Fig. S16 & S17). The gnathostome and tetrapod ancestral GKRPs inhibit their corresponding GCKs (Fig. S18 & S19). Addition of S6P does not alter their inhibitory activity (Fig. S20 & S21). Our measurements suggest that small-molecule allostery in GKRP is a mammal specific trait, which is consistent with previous comparative observations (17, 30).

Among mammals, allosteric modulation of GKRP inhibition of GCK by phosphorylated carbohydrates has been demonstrated in rabbit, rat, and human (30). Expanding upon these findings, we measured the GKRP inhibition of GCK in proteins from the wombat *Vombatus ursinus*, a marsupial. Wombat GKRP inhibited its corresponding GCK with a micromolar IC_50_ value similar to the gnathostome and tetrapod ancestral GKRPs (Fig. S22). Addition of S6P does not alter this IC_50_ value (Fig. S23). Stepwise resurrection of ancestral mammalian GKRPs revealed the emergence of small-molecule allosteric regulation. The inhibitory activity of the GKRP corresponding to the ancestor of placental mammals (M1; Fig. 3A) is insensitive to S6P (Fig. S24 & S25). Resurrection and characterization of the ancestor of the euarchontoglires (M2; Fig. 3A) — the clade containing rabbits, rodents, and primates — shows that small-molecule allostery is specific to this group, as M2 GKRP displays sensitivity to S6P (Fig. S26 & S27).

**Figure 3.**
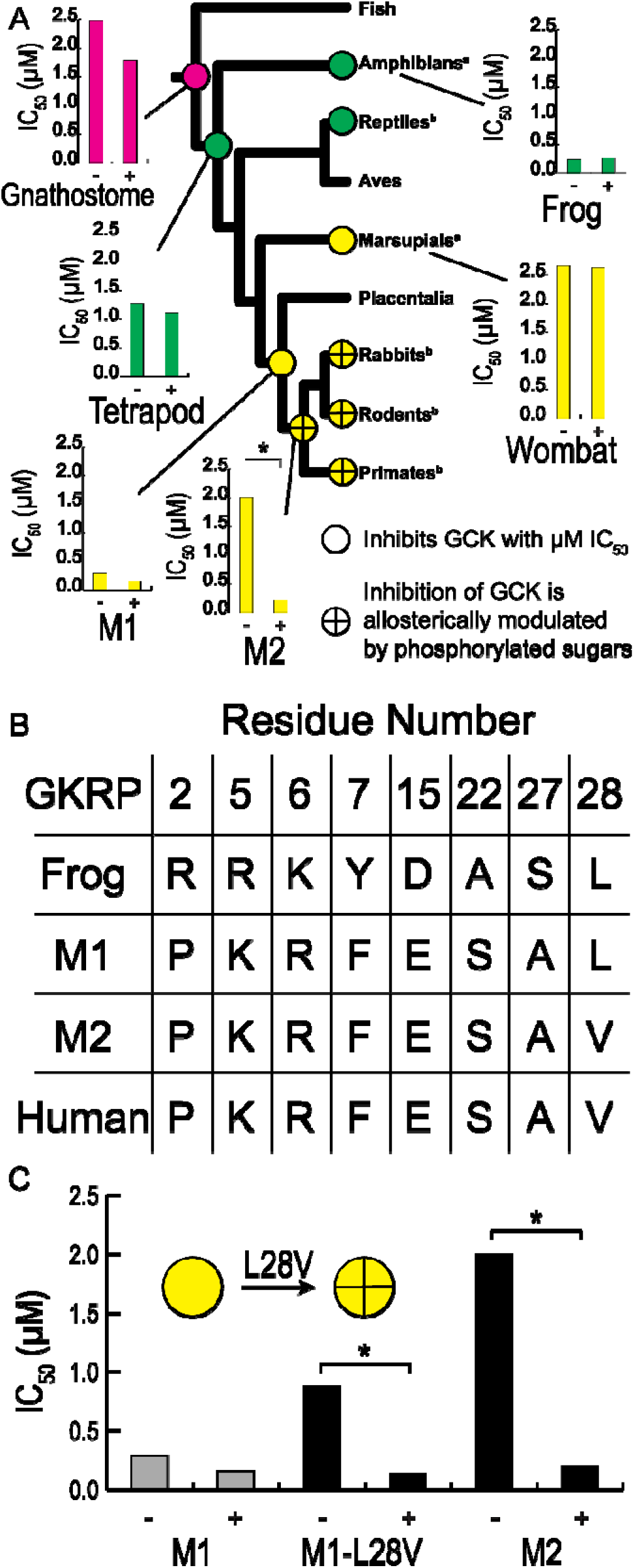
Allosteric regulation of GKRP inhibition of GCK evolved via a loss-of-function substitution in the Euarchontoglires ancestor. (*A*) Measurements of extant and extinct GKRPs’ inhibition of GCK in the absence (-) and presence (+) of sorbitol-6-phosphate (2 mM) along the vertebrate evolutionary trajectory. A functional shift in sensitivity to phosphorylated carbohydrates occurred between the ancestor of placental mammals (M1) and the ancestor of the euarchontoglires (M2). Bars represent the average of two biological replicates. * - indicates significant difference, p < 0.05. ^a^ – data from this study; ^b^ – from (30). (*B*) The N-terminus of GKRP (residues 1-30) is highly conserved among tetrapods. Positions with amino acid substitutions are shown. There is a single substitution between the M1 and M2 ancestors. (*C*) The L28V substitution in the M1 GKRP ancestor background introduces allosteric regulation by reducing the inhibition activity in the absence of phosphorylated carbohydrate while leaving activity unchanged in the presence. Bars represent the average of two biological replicates. See SI Table 2 for error estimates. * indicates significant difference, p < 0.05.

The N-terminus of GKRP (residues 1-30 in human GKRP numbering) is the structural feature that likely modulates the GCK binding interface in response to phosphorylated carbohydrate binding (31). We compared the frog, M1, M2, and human GKRP sequences to determine if there are substitutions in the N-terminus that are responsible for small-molecule allostery. Only eight residues show variation among the four GKRPs, while the rest are conserved (Fig. 3B). Of these eight, only one residue has a substitution between M1 and M2 — an L to V substitution at position 28. We introduced the M2 residue into the non-allosterically regulated M1 background. Measurement of GCK inhibition demonstrates that the resulting loss of a single methylene group via the L28V substitution is sufficient to introduce sensitivity to S6P (Fig. 3C; Fig. S28 & S29).

Counterintuitively, the L28V substitution imparts allostery through the loss of inhibitory function in the apo-state of GKRP while leaving activity in the presence of S6P unchanged (Fig. 3C). In the rat-human GKRP-GCK complex, ligand binding to GKRP kinetically stabilizes the GKRP-GCK bound state by increasing the rate constant for forming the encounter complex (*k*_on_) and by decreasing the rate constant for dissociation (*k*_off_) (32). Based on structural data, the rat GKRP N-terminus samples at least two conformations (31). In the S6P bound, inhibitory-competent state, the N-terminus is extended where it can influence residues at the GCK binding interface, likely by stabilizing them in the binding competent state (Fig. 1A). In the F1P bound, inhibitory-incompetent state, the N-terminus is folded at the interface of the LID and SIS domains, far removed from the GCK binding site. We postulate that the L28V substitution affords access to this second conformation by ordering the N-terminus in the apo-state. Martinez *et al*. produced a truncated rat GKRP missing the first twenty residues (31). This version of rat GKRP loses inhibitory activity in the presence of S6P while activity in the absence remains unchanged. Eliminating the ability to sample the two different N-terminal conformations removes allosteric modulation, providing biochemical and biophysical mechanisms for the emergence of small-molecule allostery consistent with our postulate.

## Conclusions

Our vertical approach allowed identification of a functional transition point along the GKRP evolutionary trajectory. A single loss-of-function cytosine to thymine substitution in the GKRP gene could lead to the L28V amino acid substitution, which is sufficient to cause co-optation of a pre-existing binding site for phosphorylated carbohydrates derived from a vestigial etherase active site. This suggests that the simplest genetic mechanism can enable ligand-mediated allosteric regulation of proteins. Importantly, the evolution of this regulatory novelty in GKRP did not require the addition of new genetic material. Due to its simplicity, we speculate that our simple model of the evolution of novel protein function is likely to be a general mechanism.

### Materials and Methods Synteny Analysis

We used the Ensembl (28) and Genomicus (29) databases to assess orthology and synteny in eukaryotic GKRPs. The GKRP phylogenetic tree (SI file) was estimated using the Neighbor-Joining method from a MUSCLE (33) alignment of GKRP amino acid sequences (SI file), all implemented in MEGA 7 (34).

### Activity Assays

Extant and extinct recombinant GKRPs and GCKs were produced as described previously (18). Recombinant *E. coli* MurK was expressed with a C-terminal hexahistidine tag. Recombinant *E. coli* MurQ was expressed with an N-terminal hexahistidine tag. Both were purified with Ni-NTA affinity chromatography and size-exclusion chromatography.

GKRP inhibition of GCK was measured as previously described (18). Extinct and extant GCKs were mixed with corresponding GKRPs at varying GKRP concentrations. The protein mixture was incubated for five minutes, with or without S6P, prior to initiation of the GCK reaction. GCK activity was followed by linked enzyme assay where production of glucose-6-phosphate by GCK is coupled to the reduction of NADP+ by glucose-6-phosphate dehydrogenase and measured in a spectrophotometer at 340 nm.

Etherase activity of *E. coli* MurQ and extinct and extant GKRPs was measured with a three-step linked enzyme assay (35). First, the etherase catalyzes the hydrolysis of the MurNAc-6P lactyl ether bond, producing GlcNAc-6P and D-lactate. Next, D-lactate dehydrogenase (D-LDH) catalyzes the transfer of D-lactate’s hydroxyl hydrogen to NAD^+^, producing pyruvate and NADH. In the last step, diaphorase catalyzes the reduction of p-iodonitrotetrazolium violet (INT) by NADH, recycling NADH into NAD^+^ and producing formazan, which absorbs at 500 nm.

The etherase substrate MurNAc-6P was produced by phosphorylation of commercially available N-acetylmuramate by MurK. MurNAc-6P was purified with HILIC HPLC (Fig. S30) on a Shimadzu HPLC system (Shimadzu, Kyoto, Japan) based on a published protocol (35) and its identity confirmed via mass spectrometry (Fig. S31). Various concentrations of D-lactate (134 **μ**M–900 **μ**M) were assayed in the absence of GKRP or MurQ. A standard curve for D-lactate was constructed based upon the maximum absorbance reached after 15 minutes of incubation. The data were fit to a linear equation of y = ax + b, where y is the maximum absorbance, x is the D-lactate concentration, a is the slope of the line, and b is the line’s y-intercept. This assay was repeated with 4 **μ**L of a MurNAc-6P sample that had been diluted 1:100 with sodium phosphate buffer (40 mM, pH 12.5) and incubated at 37° C overnight. The maximum absorbance obtained was used in the standard curve’s linear equation (y = 1.952x – 0.007253) to calculate the concentration of D-lactate produced, which was then used to calculate the concentration of MurNAc-6P in the original stock solution. Data points for the standard curve and the MurNAc-6P sample were collected in triplicate.

Full experimental details can be found in SI Materials and Methods.

## Acknowledgments

Research reported in this publication was supported by the National Institute of General Medical Sciences of the National Institutes of Health under Award Number R01GM133843 (B.G.M.) The content is solely the responsibility of the authors and does not necessarily represent the official views of the National Institutes of Health. Additional funding was provided by the FSU Council on Research and Creativity (B.G.M).

## References

1. G. B. Golding, A. M. Dean, The structural basis of molecular adaptation. Mol. Biol. Evol. 15, 355–369 (1998).

2. A. M. Dean, J. W. Thornton, Mechanistic approaches to the study of evolution: The functional synthesis. Nat. Rev. Genet. 8, 675–688 (2007).

3. C. Schulenburg, B. G. Miller, Enzyme Recruitment and Its Role in Metabolic Expansion. Biochemistry 53, 836–845 (2014).

4. P. A. Steindel, E. H. Chen, J. D. Wirth, D. L. Theobald, Gradual neofunctionalization in the convergent evolution of trichomonad lactate and malate dehydrogenases. Protein Science 25, 1319–1331 (2016).

5. J. W. Thornton, Evolution of vertebrate steroid receptors from an ancestral estrogen receptor by ligand exploitation and serial genome expansions. Proc. Natl. Acad. Sci. U. S. A. 98, 5671 (2001).

6. R. P. Bhattacharyya, A. Reményi, B. J. Yeh, W. A. Lim, Domains, motifs, and scaffolds: the role of modular interactions in the evolution and wiring of cell signaling circuits. Annu. Rev. Biochem. 75, 655–680 (2006).

7. A. Iorio, C. Brochier-Armanet, C. Mas, F. Sterpone, D. Madern, Protein Conformational Space at the Edge of Allostery: Turning a Nonallosteric Malate Dehydrogenase into an “Allosterized” Enzyme Using Evolution-Guided Punctual Mutations. Mol. Biol. Evol. 39 (2022).

8. A. Hadzipasic, et al., Ancient origins of allosteric activation in a Ser-Thr kinase. Science (1979). 367, 912–917 (2020).

9. S. Ohno, Evolution by Gene Duplication (Springer Berlin Heidelberg, 1970).

10. A. C. Whittington, A. J. Mason, D. R. Rokyta, A single mutation unlocks cascading exaptations in the origin of a potent pitviper neurotoxin. Mol. Biol. Evol. 35, 887–898 (2018).

11. J. Zhang, Evolution by gene duplication: an update. Trends Ecol. Evol. 18, 292–298 (2003).

12. E. Morett, P. Bork, Evolution of new protein function: recombinational enhancer Fis originated by horizontal gene transfer from the transcriptional regulator NtrC. FEBS Lett. 433, 108–112 (1998).

13. K. S. Bonham, B. E. Wolfe, R. J. Dutton, Extensive horizontal gene transfer in cheese-associated bacteria. Elife 6 (2017).

14. S. B. Van Oss, A. R. Carvunis, De novo gene birth. PLoS Genet. 15, e1008160 (2019).

15. A. Lange, et al., Structural and functional characterization of a putative de novo gene in Drosophila. Nature Communications 2021 12:1 12, 1–13 (2021).

16. M. Lynch, J. S. Conery, The evolutionary fate and consequences of duplicate genes. Science 290, 1151–5 (2000).

17. M. Veiga-da-Cunha, T. Sokolova, F. Opperdoes, E. Van Schaftingen, Evolution of vertebrate glucokinase regulatory protein from a bacterial N-acetylmuramate 6-phosphate etherase. Biochem. J. 423, 323–32 (2009).

18. S. S. Kamalaldinezabadi, et al., Evolution of Protein Regulation in the Vertebrate Glucose Sensor. bioRxiv 2026.05.05.723016 (2026). 10.64898/2026.05.05.723016.

19. F. M. Matschinsky, Glucokinase as glucose sensor and metabolic signal generator in pancreatic beta-cells and hepatocytes. Diabetes 39, 647–652 (1990).

20. T. Beck, B. G. Miller, Structural basis for regulation of human glucokinase by glucokinase regulatory protein. Biochemistry 52, 6232–6239 (2013).

21. E. Van Schaftingen, A. Vandercammen, M. Detheux, D. R. Davies, The regulatory protein of liver glucokinase. Adv. Enzyme Regul. 32, 133–148 (1992).

22. C. Shiota, J. Coffey, J. Grimsby, J. F. Grippo, M. A. Magnuson, Nuclear import of hepatic glucokinase depends upon glucokinase regulatory protein, whereas export is due to a nuclear export signal sequence in glucokinase. J. Biol. Chem. 274, 37125–37130 (1999).

23. A. Pautsch, et al., Crystal structure of glucokinase regulatory protein. Biochemistry 52, 3523–3531 (2013).

24. J. M. Choi, M. H. Seo, H. H. Kyeong, E. Kim, H. S. Kim, Molecular basis for the role of glucokinase regulatory protein as the allosteric switch for glucokinase. Proc. Natl. Acad. Sci. U. S. A. 110, 10171–10176 (2013).

25. T. Jaeger, M. Arsic, C. Mayer, Scission of the lactyl ether bond of N-acetylmuramic acid by Escherichia coli “etherase.” J. Biol. Chem. 280, 30100–30106 (2005).

26. T. Jaeger, C. Mayer, N-acetylmuramic acid 6-phosphate lyases (MurNAc etherases): role in cell wall metabolism, distribution, structure, and mechanism. Cell. Mol. Life Sci. 65, 928–939 (2008).

27. T. Hadi, S. Hazra, M. E. Tanner, J. S. Blanchard, Structure of MurNAc 6-phosphate hydrolase (MurQ) from Haemophilus influenzae with a bound inhibitor. Biochemistry 52, 9358–9366 (2013).

28. A. D. Yates, et al., Ensembl Genomes 2022: an expanding genome resource for non-vertebrates. Nucleic Acids Res. 50, D996–D1003 (2022).

29. N. T. T. Nguyen, P. Vincens, H. R. Crollius, A. Louis, Genomicus 2018: karyotype evolutionary trees and on-the-fly synteny computing. Nucleic Acids Res. 46, D816–D822 (2018).

30. A. Vandercammen, E. Van Schaftingent, Species and tissue distribution of the regulatory protein of glucokinase. 294, 551–556 (1993).

31. J. A. Martinez, Q. Xiao, A. Zakarian, B. G. Miller, Antidiabetic Disruptors of the Glucokinase-Glucokinase Regulatory Protein Complex Reorganize a Coulombic Interface. Biochemistry 56, 3150–3157 (2017).

32. A. K. Casey, B. G. Miller, Kinetic Basis of Carbohydrate-Mediated Inhibition of Human Glucokinase by the Glucokinase Regulatory Protein. Biochemistry 55, 2899–2902 (2016).

33. R. C. Edgar, MUSCLE: multiple sequence alignment with high accuracy and high throughput. Nucleic Acids Res. 32, 1792 (2004).

34. S. Kumar, G. Stecher, K. Tamura, MEGA7: Molecular Evolutionary Genetics Analysis Version 7.0 for Bigger Datasets. Mol. Biol. Evol. 33, 1870–1874 (2016).

35. S. Unsleber, M. Borisova, C. Mayer, Enzymatic synthesis and semi-preparative isolation of N-acetylmuramic acid 6-phosphate. Carbohydr. Res. 445, 98–103 (2017).

