## Supplementary Text for "Structural Co-optation and Loss-of-function Underlie the Evolution of Regulatory Novelty in the Glucokinase Regulatory Protein"

**This PDF file includes:**

Material and Methods

Figures S1 to S32

Tables S1 and S2

SI References

**Other supporting materials for this manuscript include the following:**

Alignment of GKRP sequences

Protein sequences used in this study

Supporting Information Text

**Materials and Methods**

**Recombinant protein production**

Recombinant GCKs and GKRPs were produced as described previously (1).

Recombinant MurK was produced as a C-terminal His6-tagged polypeptide in the pET-28a(+) vector in E. coli BL21 (DE3) cells. A culture was inoculated to an OD600 of 0.02 in LB supplemented with kanamycin (50 μg/mL). Cells were shaken at 250 rpm, 37 °C until the OD600 reached 0.70, at which point IPTG (0.1 mM) was added to induce gene expression. Growth continued for 3 h, then the cells were harvested by centrifugation for 10 min at 6,000 × g and 4 °C. Wet cell pellets were weighed and stored at -20 °C until ready for use.

Frozen MurK cell pellets were resuspended in cold MurK loading buffer (5 mL/g of pellet) containing sodium phosphate (20 mM, pH 7.5), NaCl (500 mM), imidazole (20 mM), and DTT (10 mM). The cells were lysed via sonication with a Branson Sonifier S-250A (Branson Ultrasonics). Cells were subjected to 10 s of pulses with a 50% duty cycle at 50% output intensity, followed by 50 s of recovery time. This was repeated 20 times. Crude cell lysates were centrifuged at 25,000 × g and 4 °C for 30 min, and the cleared cell lysates were loaded onto a 5 mL HisTrap FF affinity column (Cytiva) equilibrated with cold MurK loading buffer. The column was washed with 10 column volumes of cold MurK loading buffer, and protein was eluted with 5 column volumes of cold MurK elution buffer (loading buffer supplemented with 100 mM imidazole). MurK eluate was dialyzed overnight at 4 °C against 1 L of MurK SEC buffer containing sodium phosphate (20 mM, pH 7.5), NaCl (500 mM), and DTT (10 mM). After dialysis, the sample was concentrated to 1 mL using an Amicon Ultra-15 Centrifugal Filter Unit with a MWCO of 10 kDa (MilliporeSigma) and injected onto a Superdex 200 Increase 10/300 column (Cytiva), pre-equilibrated in MurK SEC buffer. The SEC column was run at a flow rate of 0.15 mL/min, and fractions containing the highest A280 values were pooled and retained for characterization. Chemicals for purification were purchased from Thermo Fisher Scientific, Sigma-Aldrich, and Gold Biotechnology.

Recombinant E. coli MurQ was produced as an N-terminal His6-tagged polypeptide from the ASKA collection40 in the pCA24N vector in E. coli BL21 (DE3) cells. A culture was inoculated to an OD600 of 0.005 in M9 minimal media supplemented with chloramphenicol (30 μg/mL). M9 minimal media contained sodium phosphate (423 mM, pH 8.2), potassium phosphate (222 mM, pH 8.2), NaCl (86 mM), NH4Cl (52 mM), glucose (1% (w/v)), thiamine (83 μM), MgCl2 (1 mM), FeSO4 (10 μM), FeCl3 (0.6 μM), ZnCl2 (0.07 μM), CoCl2 (0.05 μM), Na2MoO4 (0.05 μM), CuSO4 (7.6 μM), H3CO3 (7.9 mM), HCl (0.12 μM), L-alanine (0.2 g/L), L-arginine (0.2 g/L), L-aspartic acid (0.2 g/L), L-asparagine monohydrate (0.2 g/L), L-glutamine (0.2 g/L), L-glutamic acid (0.2 g/L), L-glycine (0.2 g/L), L-histidine (0.2 g/L), L-lysine (0.2 g/L), L-leucine (0.2 g/L), L-isoleucine (0.2 g/L), L-serine (0.2 g/L), L-threonine (0.2 g/L), L-tryptophan (0.2 g/L), L-proline (0.2 g/L), L-valine (0.2 g/L), L-methionine (0.2 g/L), and L-phenylalanine (0.2 g/L). Cells were shaken at 250 rpm, 37 °C until the OD600 reached 0.60, at which point, the temperature was decreased to 18° C and IPTG (0.4 mM) was added to induce gene expression. Growth continued for 60 h, then the cells were harvested by centrifugation for 10 min at 6,000 × g and 4 °C. Wet cell pellets were weighed and stored at -20 °C until ready for use.

Frozen MurQ cell pellets were resuspended in cold MurQ loading buffer (5 mL/g of pellet) containing HEPES (50 mM, pH 7.4), NaCl (300 mM), imidazole (25 mM), glycerol (10% (w/v)), and DTT (1 mM). The cells were lysed three times using a French press (Thermo Fisher Scientific). Crude cell lysates were centrifuged at 25,000 × g and 4 °C for 30 min, and the cleared cell lysates were loaded onto a 5 mL HisTrap FF affinity column (Cytiva) equilibrated with cold MurQ loading buffer. The column was washed with 10 column volumes of cold MurQ loading buffer, and protein was eluted with 5 column volumes of cold MurQ elution buffer (loading buffer supplemented with 250 mM imidazole). MurQ eluate was dialyzed overnight at 4 °C against 1 L of MurQ SEC buffer containing HEPES (50 mM, pH 7.4), KCl (25 mM), and DTT (1 mM). After dialysis, the sample was concentrated to 1 mL using an Amicon Ultra-15 Centrifugal Filter Unit with a MWCO of 10 kDa (MilliporeSigma) and injected onto a Superdex 200 Increase 10/300 column (Cytiva), pre-equilibrated in MurQ SEC buffer. The SEC column was run at a flow rate of 0.15 mL/min, and fractions containing the highest A280 values were pooled and retained for characterization. Chemicals for purification were purchased from Thermo Fisher Scientific, Sigma-Aldrich, and Gold Biotechnology. The QuikChange II kit (Agilent, Santa Clara, CA) was used for site-directed mutagenesis.

**Steady-state kinetics analyses of GCKs’ inhibition by GKRPs**

For steady-state kinetics analyses, GCKs were used at a final concentration of 25–100 nM, such that the uninhibited rate of glucose conversion corresponded to a rate of glucose 6-phosphate (G6P) production of ~540 nM per second. Ancestral GCKs were mixed with ancestral GKRPs at various GKRP concentrations (0–100 μM), and the proteins were incubated in GKRP SEC buffer supplemented with 10 mM DTT and, if applicable, sorbitol 6-phosphate (S6P) (final assay concentration of 2 mM) for 5 min at 25 °C to allow binding to reach equilibrium. Incubation took place in a 100 μL quartz cuvette to minimize the risk of sample loss during transfer. Once incubation was concluded, 1 unit of glucose 6-phosphate dehydrogenase (G6PDH) from Leuconostoc mesenteroides (Sigma-Aldrich) was added to the reaction mixture, followed by a master mix containing HEPES (250 mM, pH 7.1), KCl (25 mM), NADP+ (0.5 mM), DTT (10 mM), MgCl2 (6 mM), and glucose. The final concentration of glucose was 5 mM when working with vGCK, gGCK, or tGCK, and 50 μM when working with cGCK, due to cGCK’s lower K0.5 value for glucose. The reaction mixtures were incubated for another 3 min prior to initiating the reaction with ATP (5 mM). This procedure measures GCK’s activity by coupling the production of G6P to the reduction of NADP+ by G6PDH. When mixed with a GKRP, the GCK may be inhibited, and its rate of glucose conversion decreases. The rate of glucose conversion was plotted as a function of GKRP concentration and the IC_50_ values were calculated using GraphPad PRISM (Dotmatics, USA) by fitting the data to a sigmoidal ligand dose-response curve,

$$y=\frac{(max - min)}{1+{10^}^{(Log{IC}_{50}-X)}}$$

where y = the rate of G6P production, x = the concentration of GKRP, max = the uninhibited rate of G6P production, and min = the rate of G6P production when GCK is saturated with GKRP) . Data that could not be fit to a sigmoidal ligand dose-response curve with an R2 value of at least 0.9 were fit to a linear equation instead to demonstrate linearity (y = ax+b, where y = the rate of G6P production, x = the concentration of GKRP, a = the slope of the line, and b = the uninhibited rate of G6P production), and were interpreted as having an IC_50_ >1500 μM. This value represents the minimum IC_50_ possible if inhibition began at the highest concentration of GKRP used in the assay.

**MurNAc-6P synthesis and purification**

N-acetylmuramate (MurNAc) was purchased from Millipore Sigma and phosphorylated to MurNAc-6P using N-acetylmuramate kinase (MurK) from Clostridium acetobutylicum, produced as described above. To synthesize MurNAc-6P, the following were mixed into 500 μL aliquots contained within a 1.5 mL tube: Tris-HCl (50 mM, pH 8.0), MgCl2 (10 mM), MurNAc (50 mM), ATP (67 mM, pH 6.7), DTT (10 mM), SEC-purified MurK (548 nM), and water. The solution was incubated at 37° C overnight, then put on ice.

MurNAc-6P was separated from the rest of the reaction mixture with a protocol adapted from (2). 20 μL of the MurNAc-6P reaction was injected onto a 4.6x150 mm Atlantis Silica (3 μm) HILIC column (Waters, Milford MA) in an HPLC system (Shimadzu, Kyoto, Japan) with a column heating chamber kept at 37 °C and a flow rate of 2.0 mL/min. The column was first washed with a mixture of 85% buffer A (100% acetonitrile) and 15% buffer B (0.1% (v/v) formic acid, 0.05% (w/v) ammonium formate, pH 3.2) for 5 min, followed by a 5 min gradient from 15% to 21% buffer B. Buffer B was kept at 21% for 10 min, followed by 5 min of washing with 100% buffer B and 5 min of re-equilibration with 15% buffer B. Absorbance was measured at 205 nm and 245 nm. MurNAc-6P and unreacted MurNAc only absorb at 205 nm, while ATP and ADP absorb at both 205 and 245 nm. The separated MurNAc-6P was rotary evaporated to remove solvent, then it was resuspended in water and stored at -20°C. The MurNAc-6P sample was assessed for purity using an Agilent 6230 TOF-MS (Agilent) in positive ion mode.

**Determining the concentration of MurNAc-6P**

Various concentrations of D-lactate (134 μM–900 μM) were assayed in the absence of GKRP or MurQ. A standard curve for D-lactate was constructed based upon the maximum absorbance reached after 15 minutes of incubation. The data were fit to a linear equation of y = ax + b, where y is the maximum absorbance, x is the D-lactate concentration, a is the slope of the line, and b is the line’s y-intercept. This assay was repeated with 4 μL of a MurNAc-6P sample that had been diluted 1:100 with sodium phosphate buffer (40 mM, pH 12.5) and incubated at 37° C overnight. The maximum absorbance obtained was used in the standard curve’s linear equation (y = 1.952x – 0.007253) to calculate the concentration of D-lactate produced, which was then used to calculate the concentration of MurNAc-6P in the original stock solution. Data points for the standard curve and the MurNAc-6P sample were collected in triplicate.

**Steady-state kinetics analyses of GKRPs’ etherase activities**

The etherase activities of E. coli MurQ and GKRPs were measured with a linked enzyme kinetics assay comprising three enzymatic steps. First, the protein of interest catalyzes the hydrolysis of the MurNAc-6P lactyl ether bond, producing GlcNAc-6P and D-lactate. Next, D-lactate dehydrogenase (D-LDH) catalyzes the transfer of D-lactate’s hydroxyl hydrogen to NAD+, producing pyruvate and NADH. In the last step, diaphorase catalyzes the reduction of p-iodonitrotetrazolium violet (INT) by NADH, recycling NADH into NAD+ and producing formazan, which absorbs at 500 nm.

A master mix was prepared containing Tris buffer (60 mM, pH 8.0), INT (6.5 mM), NAD+ (5 mM), bovine serum albumin (BSA) (1% (w/v)) (Millipore Sigma), diaphorase from Clostridium kluyveri (0.02 U/μL) (Millipore Sigma), and D-LDH from Lactobacillus leichmannii (0.24 U/μL) (Millipore Sigma). The D-LDH and diaphorase stock solutions were kept in potassium phosphate buffer (100 mM, pH 7.0) supplemented with BSA (1% (w/v)). The master mix was mixed with varying concentrations of MurNAc-6P in a clear bottom 96-well microplate (Thermo Fisher Scientific), and the reaction was initiated by adding the protein of interest, bringing the solution to a final volume of 20 μL. The protein concentration was chosen by assaying varying protein concentrations with 625 μM MurNAc-6P, such that a reaction rate of at least 0.002 AU/s was observed. Each measurement at a particular MurNAc-6P concentration was performed in triplicate.

After reaction initiation, the microplate was inserted into a SpectraMax iD5 microplate reader (Molecular Devices, San Jose, CA) kept at 30 °C, and the increase in absorbance at 500 nm was monitored for 20 minutes. Least-squares analysis was used to calculate the highest observable rate sustained over a period of at least one minute, representing the reaction rate for that data point. Observed rates were plotted and fitted via non-linear regression with the Michaelis-Menten equation or the standard equation for reversible competitive inhibition in the case of substrate inhibition in GraphPad PRISM (Dotmatics, USA).

Figures

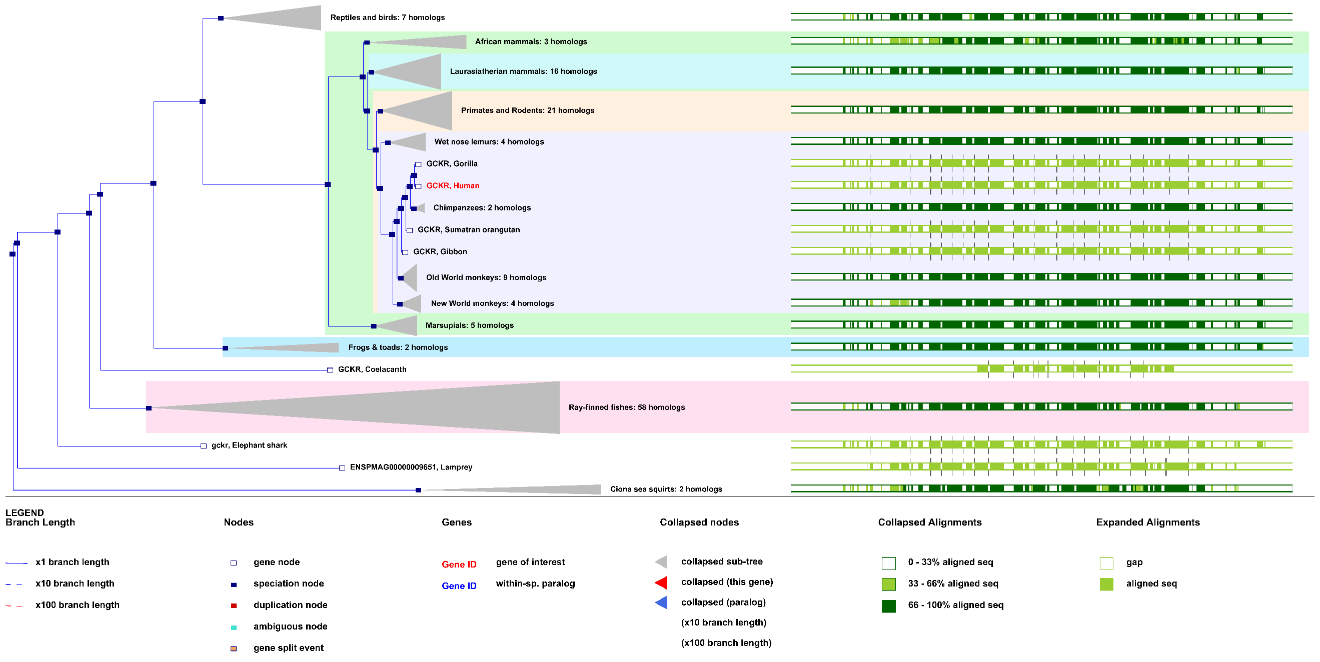

Fig. S1. Ensembl *gkrp* gene tree (release 114, May 2025) showing that among tunicates and chordates, GKRP exists as a single ortholog.

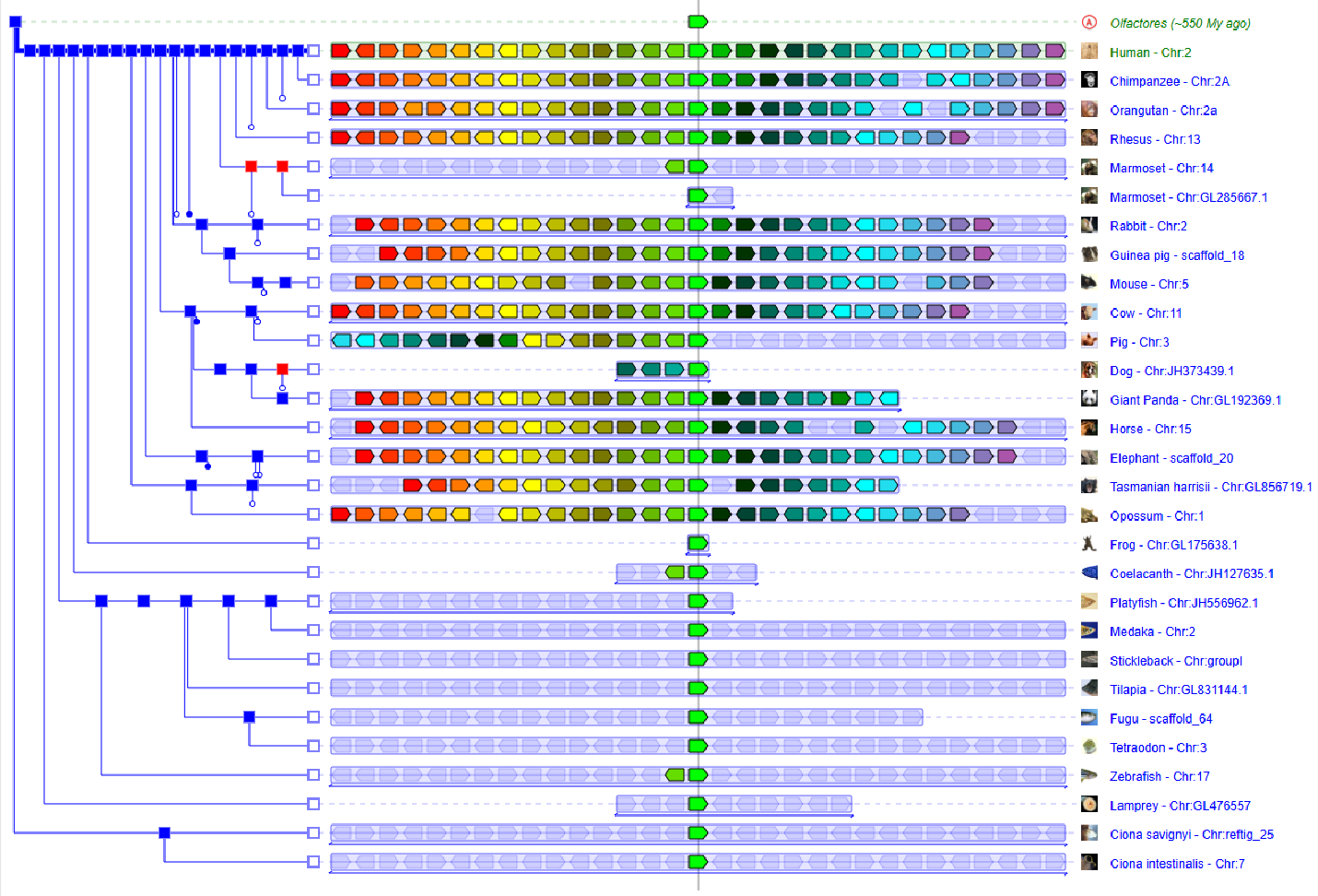

**Fig. S2**. Genomicus output showing orthology in GKRPs between tunicates and chordates, as well as showing syntenic orthology among the jawed vertebrates.

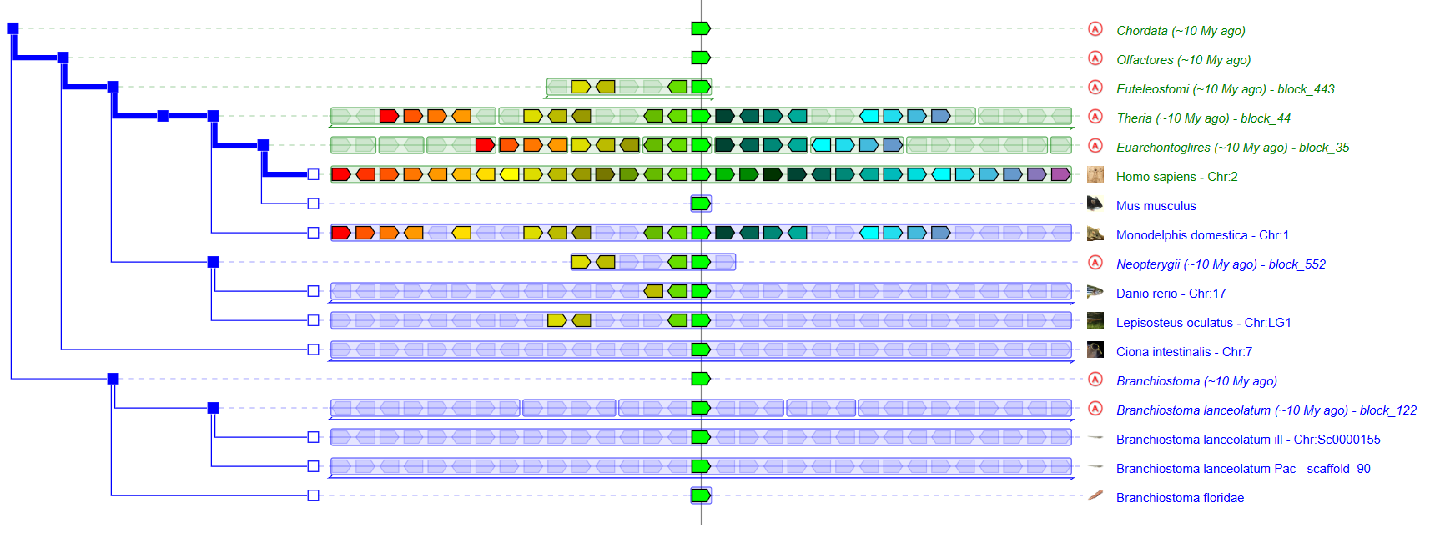

**Fig. S3**. Genomicus output showing GKRP orthologs among chordates, tunicates, and cephalochordates. Additionally, this shows further evidence of syntenic orthology of GKRPs in jawed vertebrates.

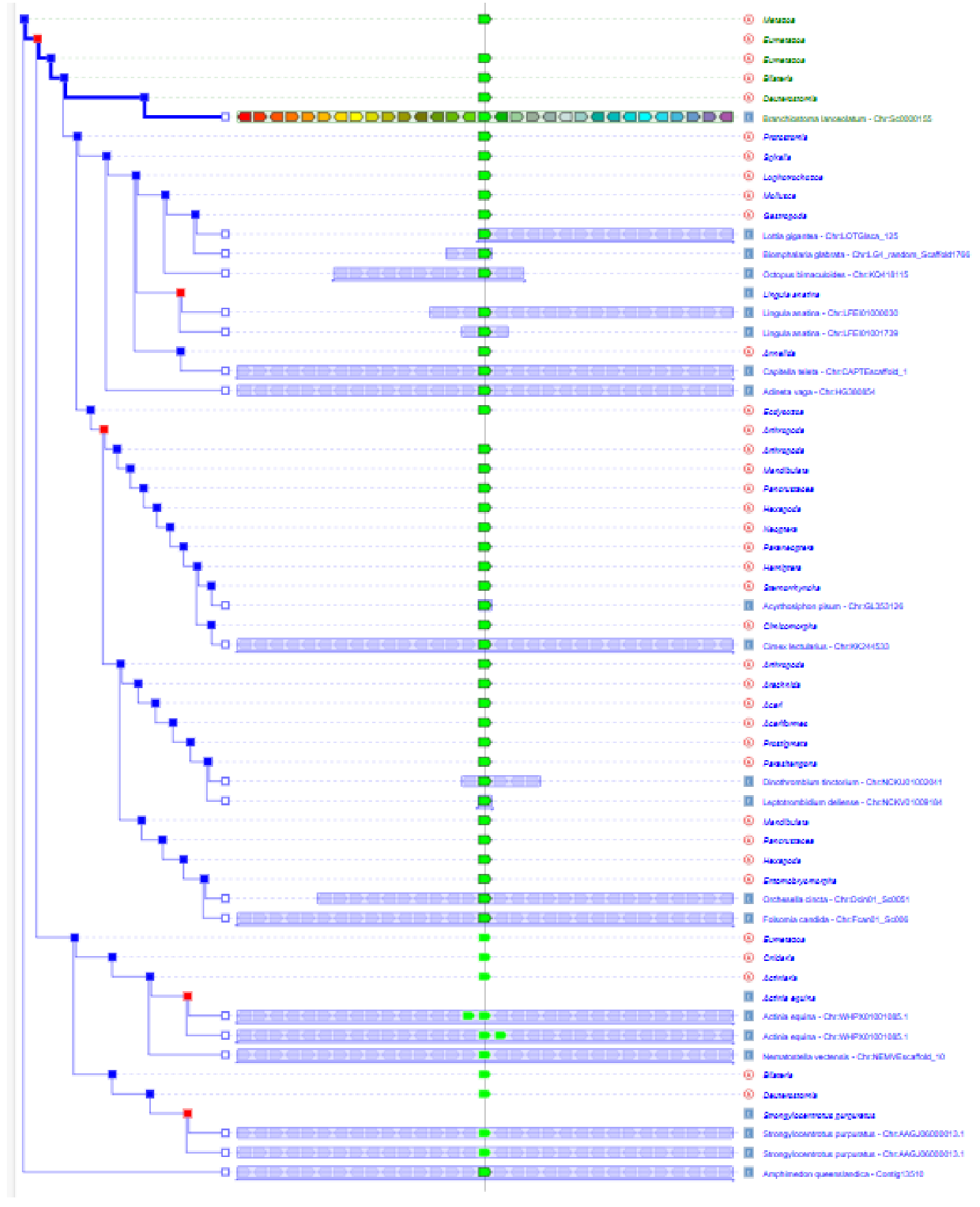

**Fig. S4**. Genomicus output showing GKRP orthology among non-chordate metazoan. The syntenic relationships are unclear. This does show the one-to-one-to-one mapping among *Branchiostoma*, *Ciona*, and *Amphimedon* of GKRP orthology.

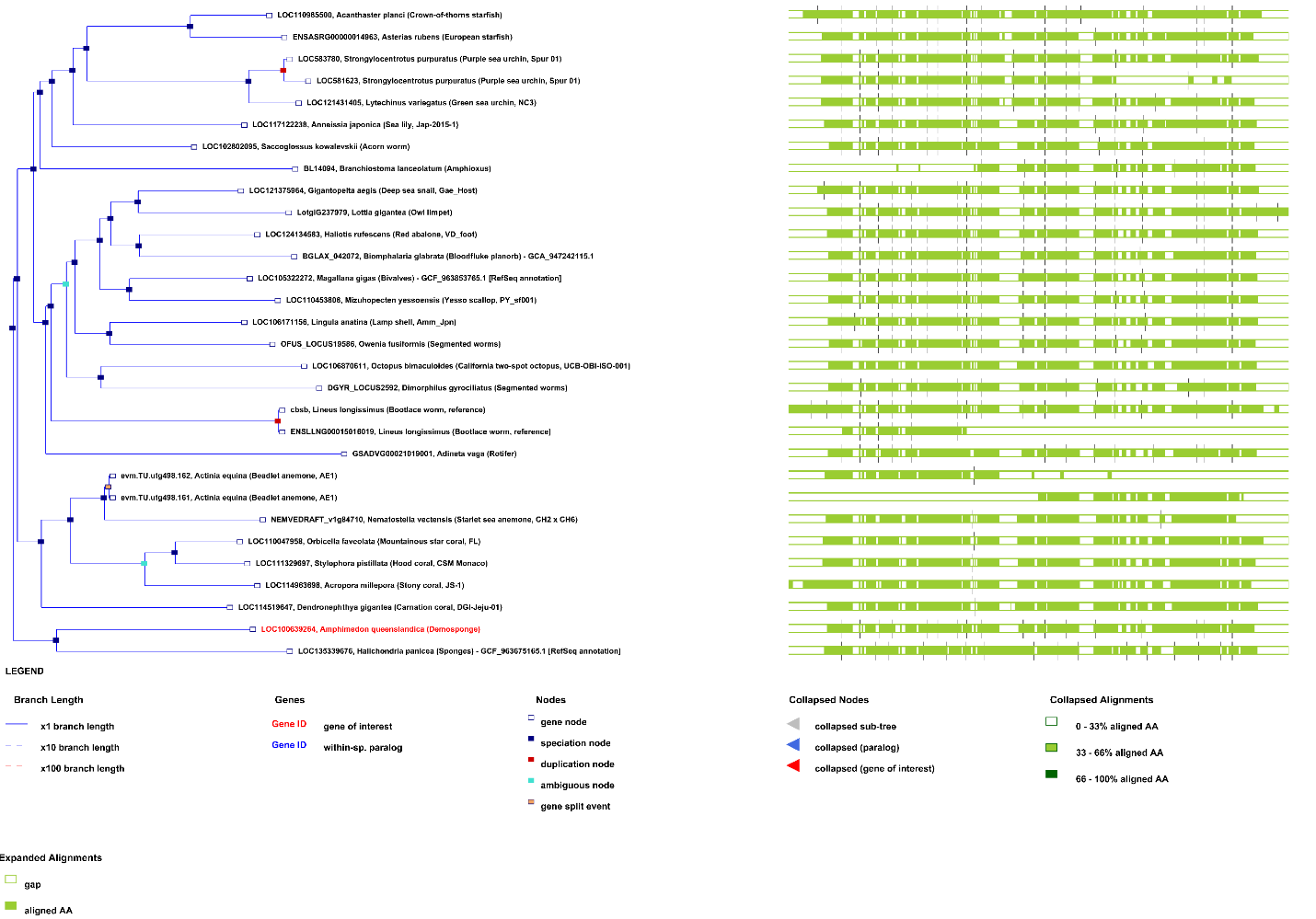

**Fig. S5**. Ensembl Metazoa *gkrp* gene tree (release 61, May 2025) showing that among metazoa, GKRP exists as a single ortholog, except for two lineage specific duplications (*Strongylocentrotus purpuratus* and *Actinia equina*), that were likely unequal based on the exon alignments. The GKRP from *Branchiostoma lanceolatum* links this tree to the chordate tree (Fig. S1), showing the orthology of GKRPs among all Metazoa including chordates.

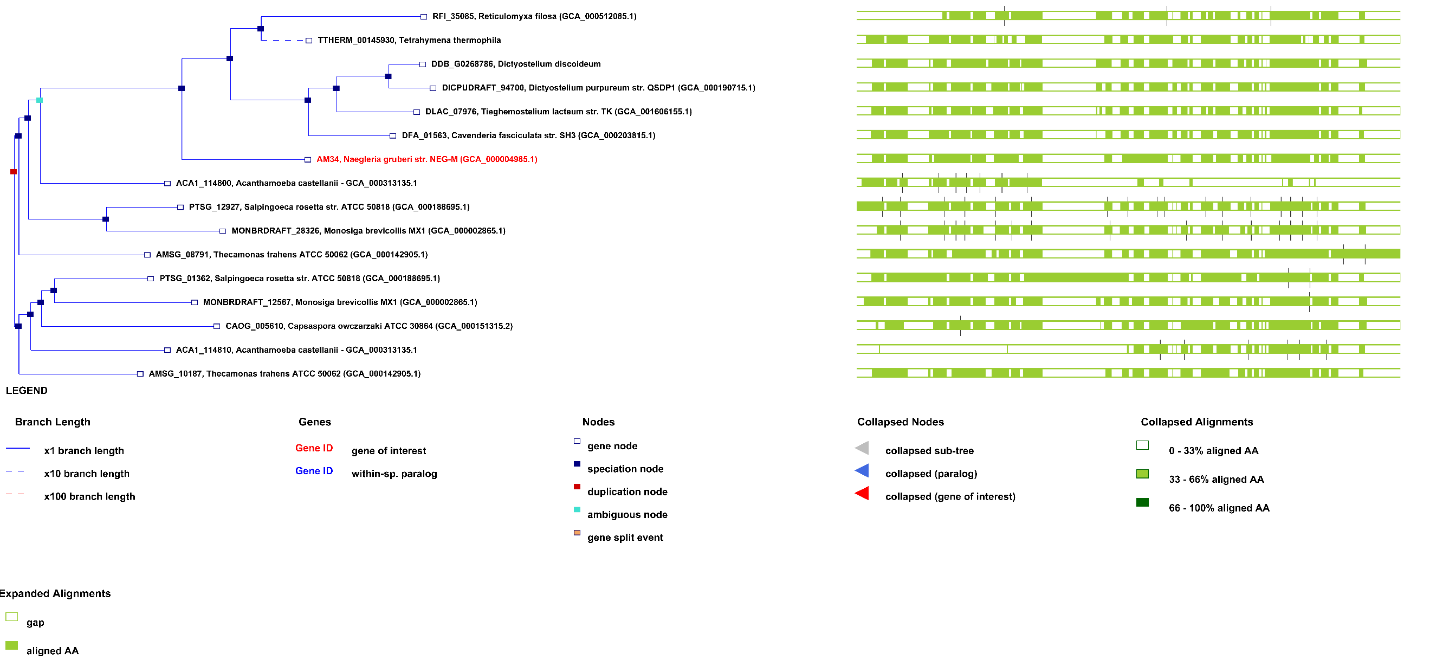

**Fig. S6**. Ensembl Protists *gkrp* gene tree (release 61, May 2025) showing a single gene duplication event. Our phylogenetic analysis (Fig. 1D) suggests that there were independent *gkrp* loses in *Capsaspora* and the Metazoan ancestor.

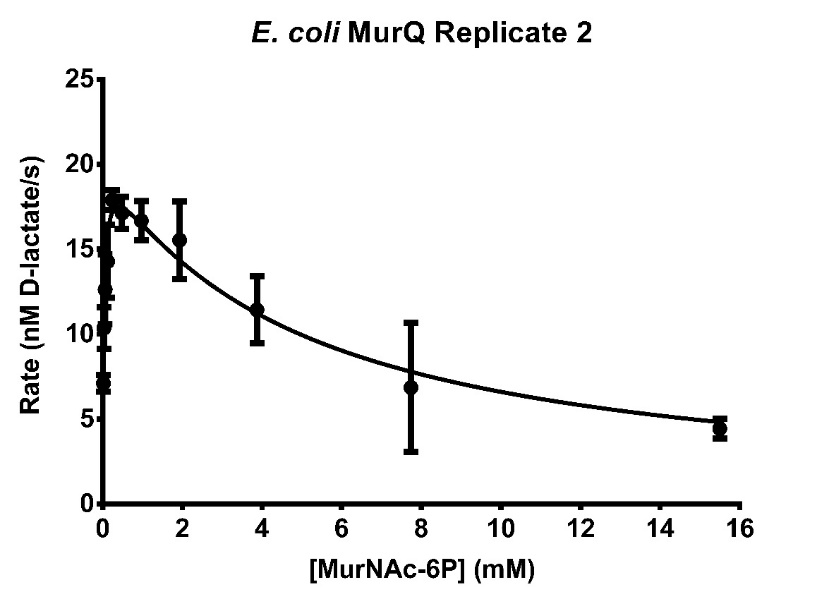

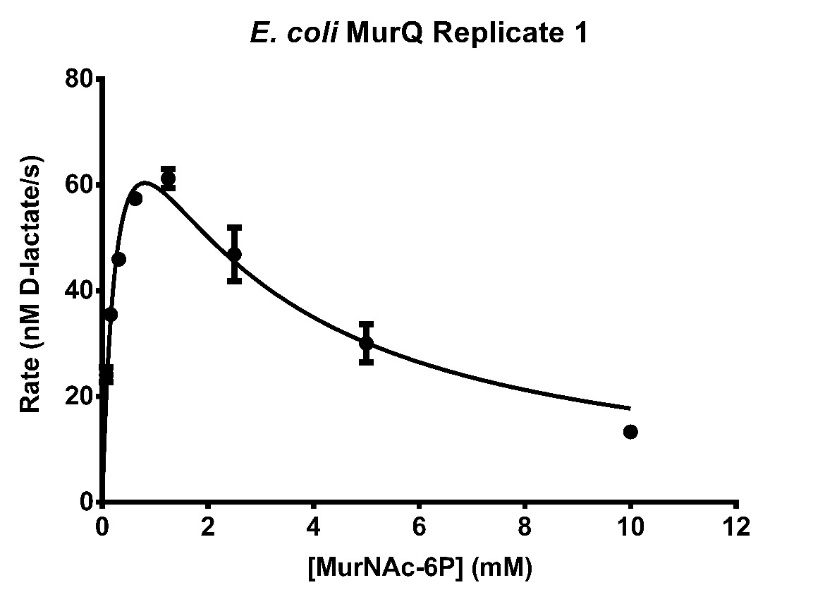

**Fig. S7:** Steady-state kinetics assay of *E. coli* MurQ’s etherase activity with varying concentrations of MurNAc-6P. Each data point represents the average of triplicate measurements of the highest rate of D-lactate production observed over at least one minute of data collection.

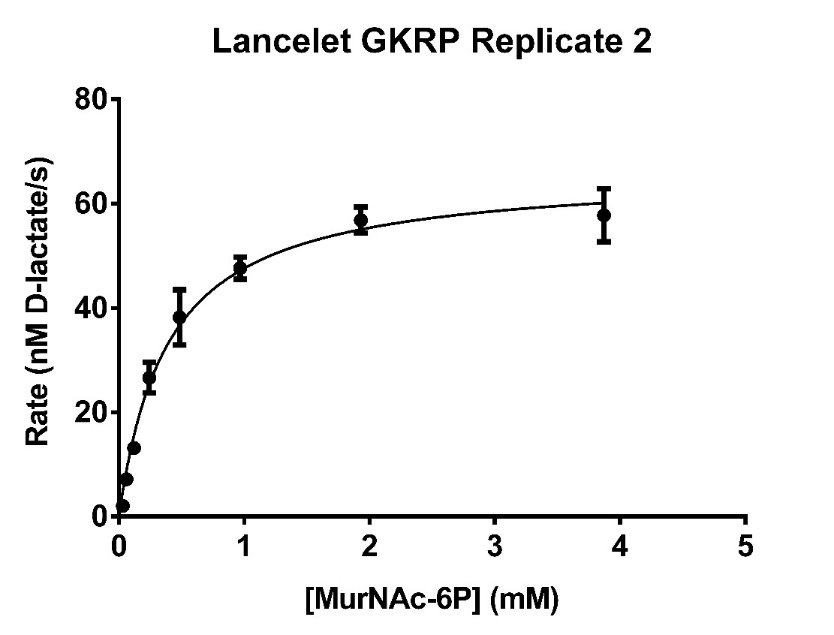

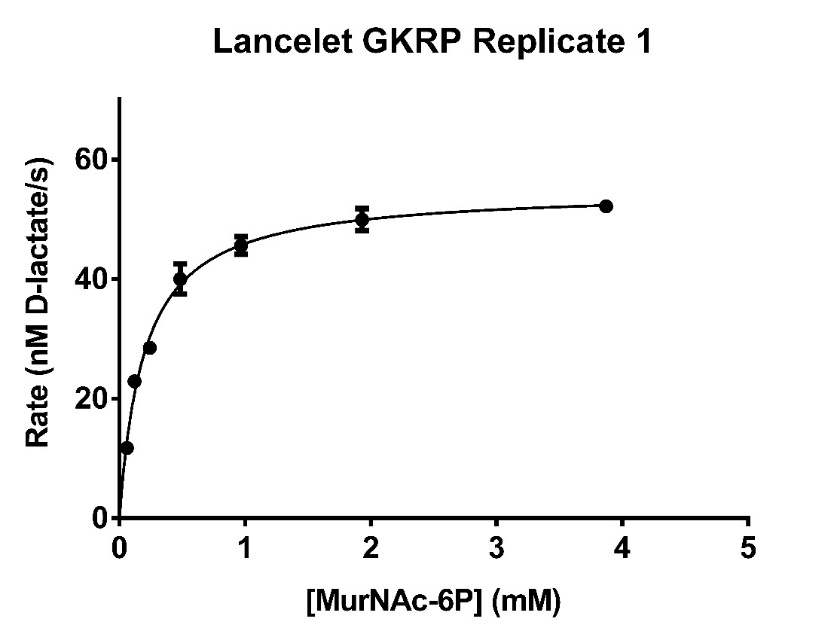

**Fig. S8:** Steady-state kinetics assay of Lancelet GKRP’s etherase activity with varying concentrations of MurNAc-6P. Each data point represents the average of triplicate measurements of the highest rate of D-lactate production observed over at least one minute of data collection.

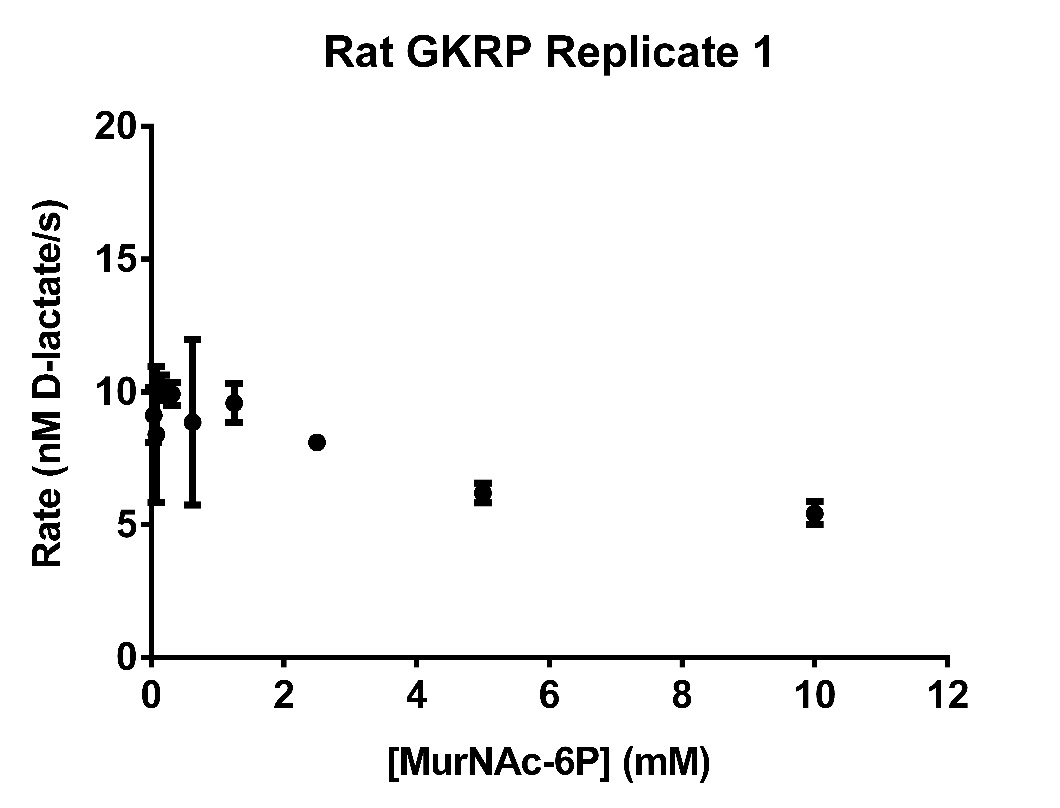

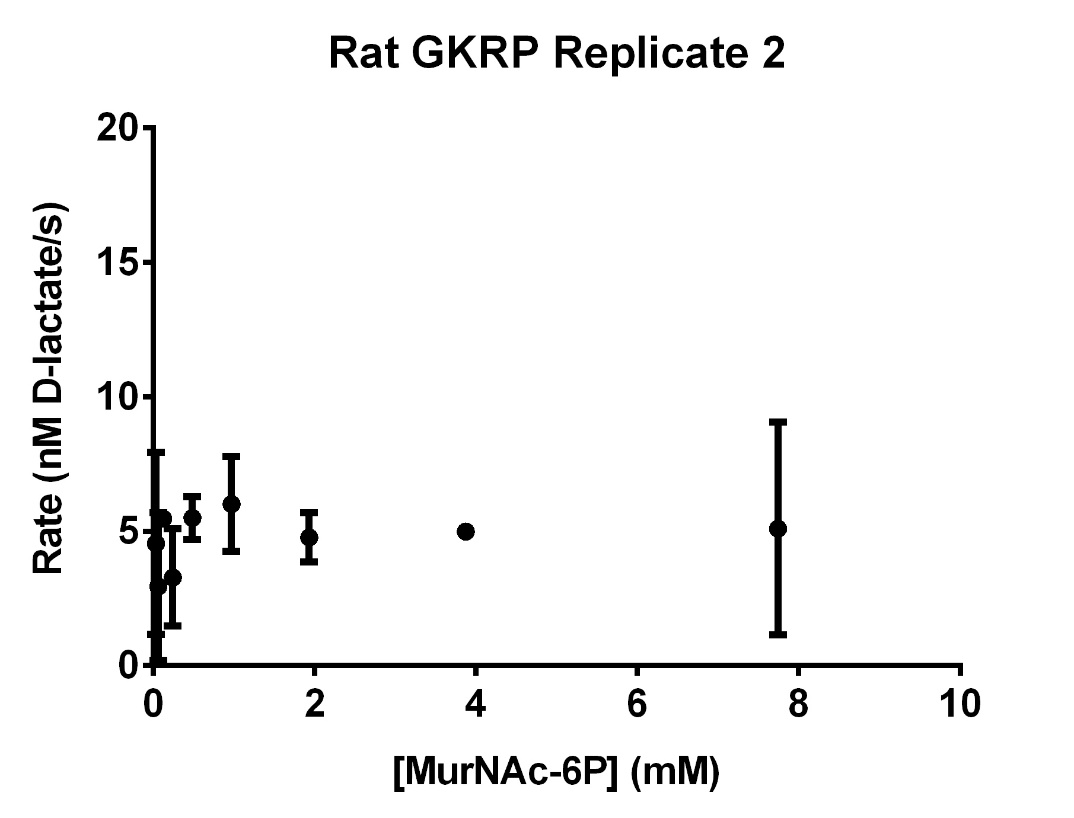

**Fig. S9:** Steady-state kinetics assay of Rat GKRP’s etherase activity with varying concentrations of MurNAc-6P. Each data point represents the average of triplicate measurements of the highest rate of D-lactate production observed over at least one minute of data collection.

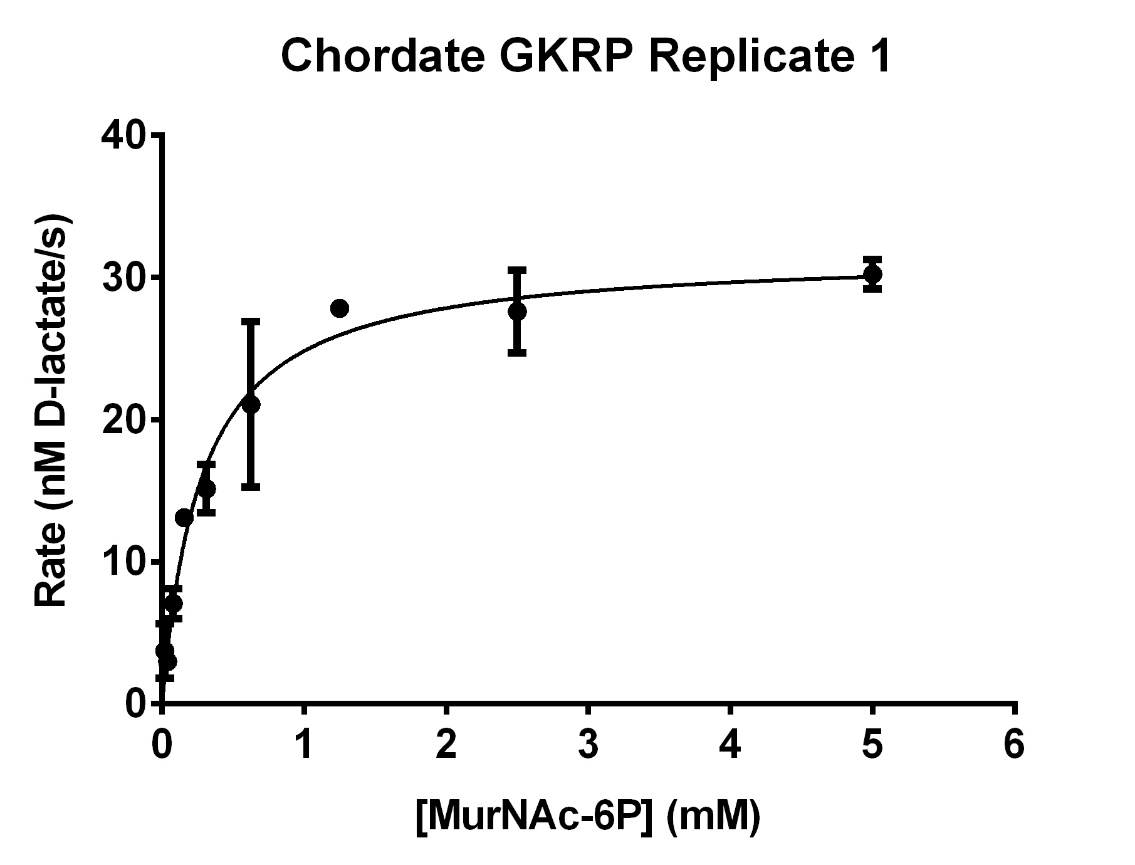

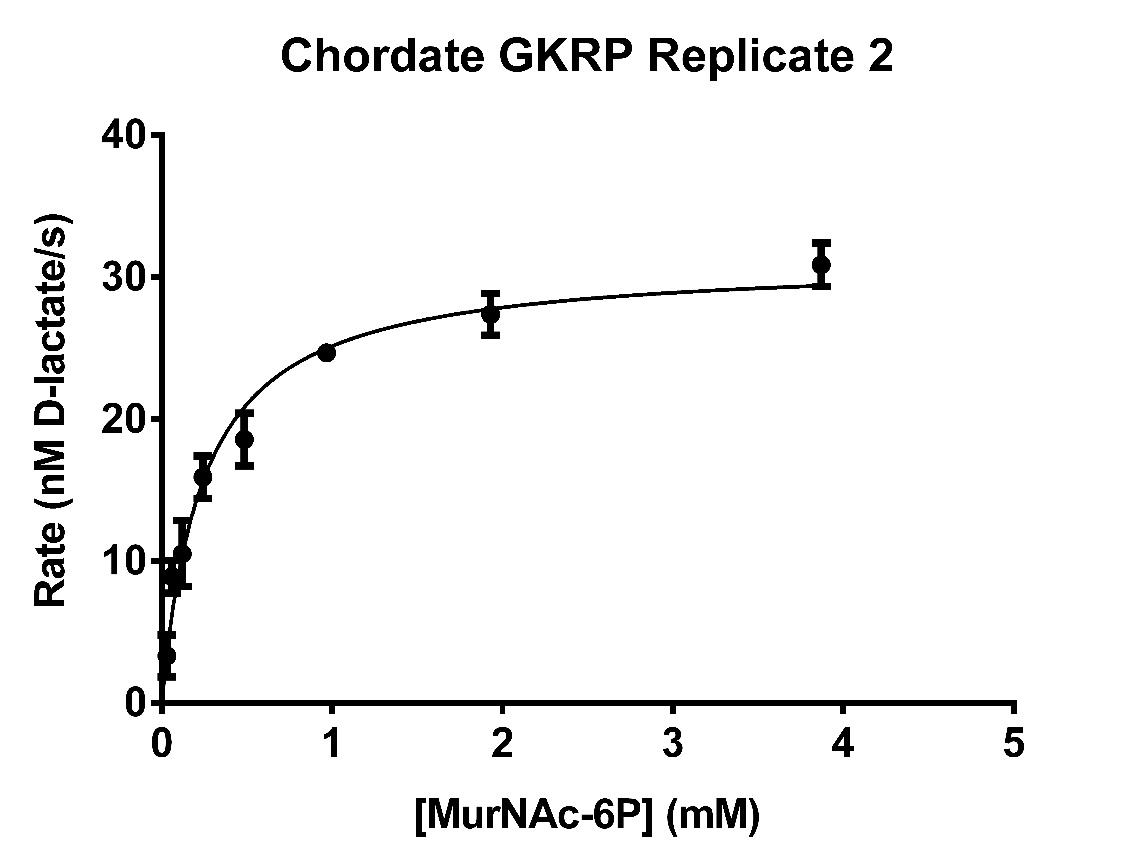

**Fig. S10:** Steady-state kinetics assay of Chordate GKRP’s etherase activity with varying concentrations of MurNAc-6P. Each data point represents the average of triplicate measurements of the highest rate of D-lactate production observed over at least one minute of data collection.

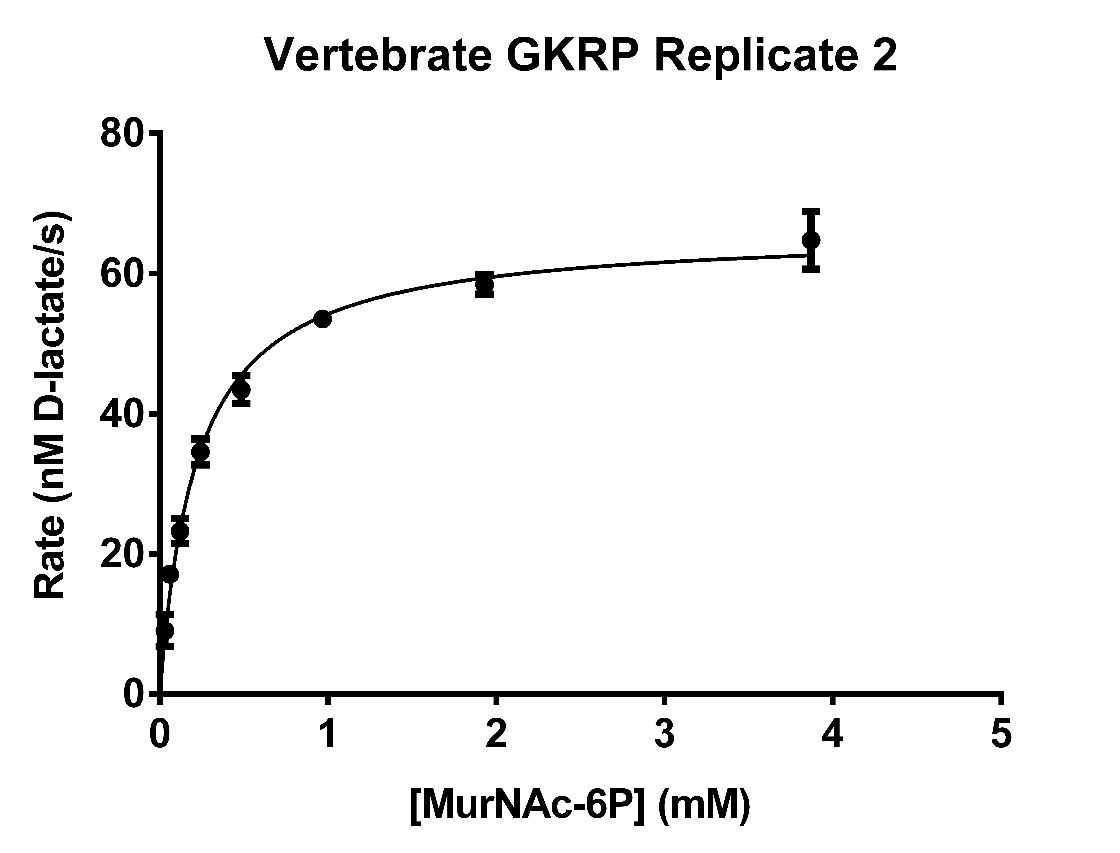

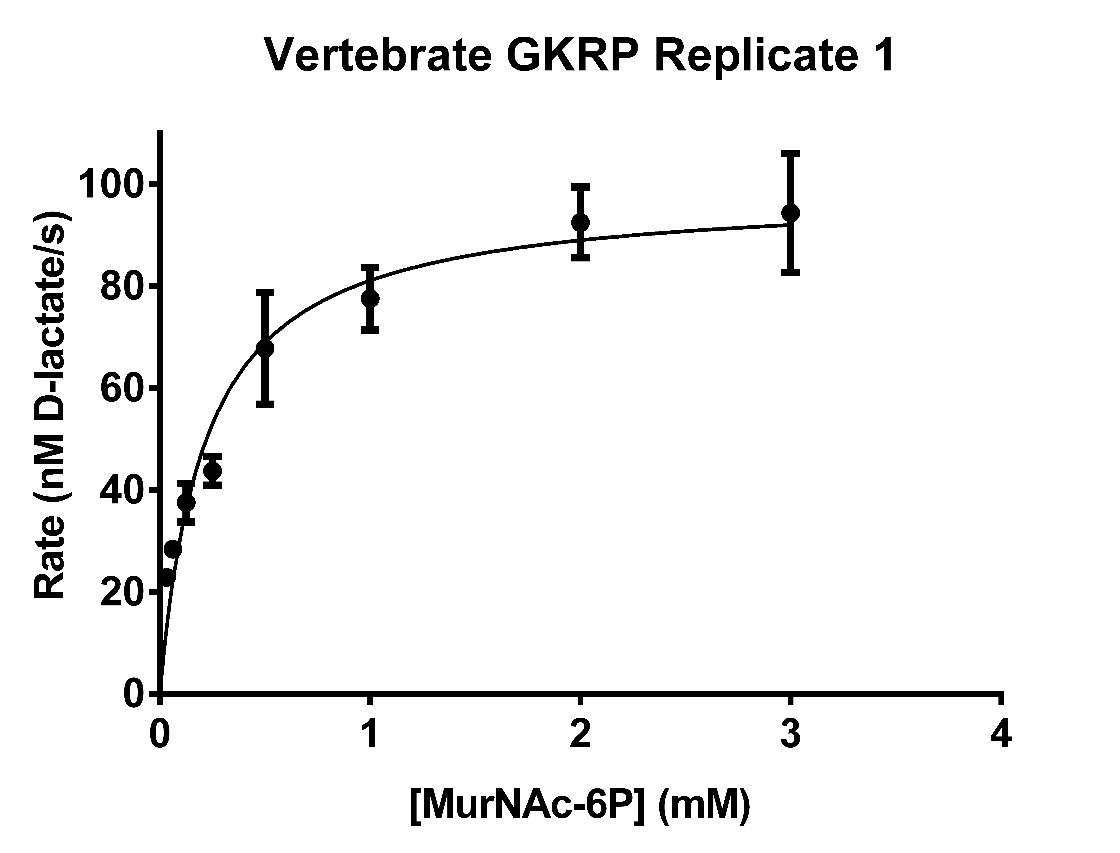

**Fig. S11:** Steady-state kinetics assay of Vertebrate GKRP’s etherase activity with varying concentrations of MurNAc-6P. Each data point represents the average of triplicate measurements of the highest rate of D-lactate production observed over at least one minute of data collection.

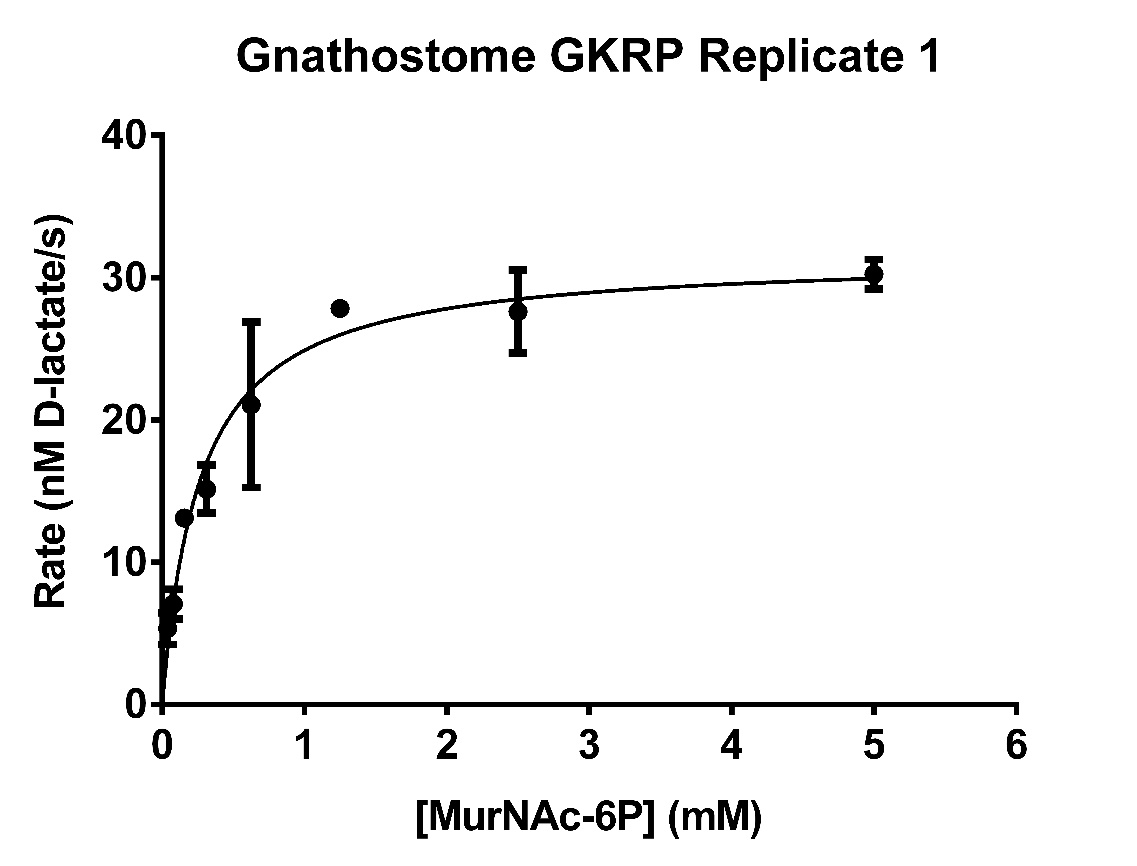

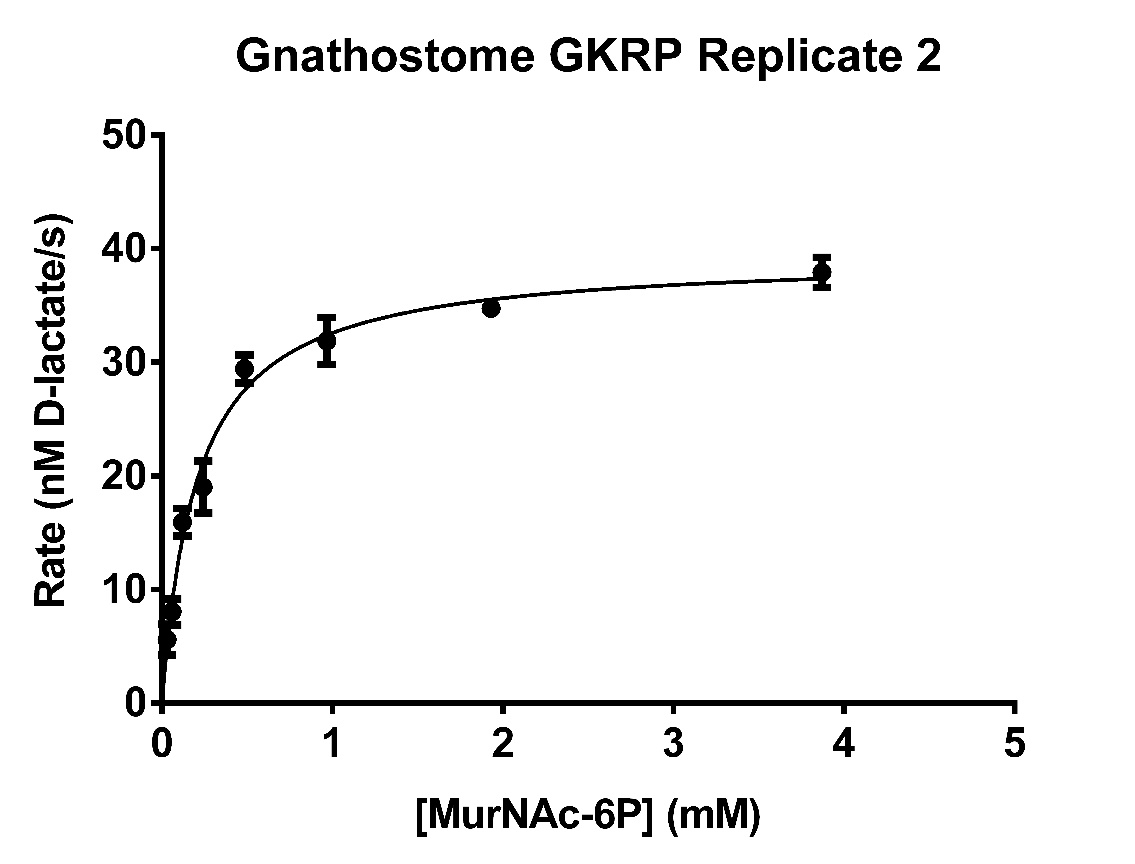

**Fig. S12:** Steady-state kinetics assay of Gnathostome GKRP’s etherase activity with varying concentrations of MurNAc-6P. Each data point represents the average of triplicate measurements of the highest rate of D-lactate production observed over at least one minute of data collection.

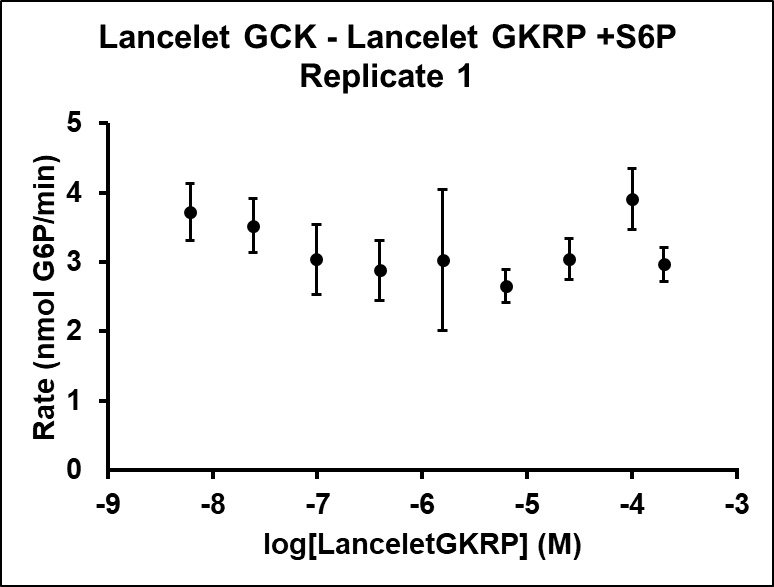

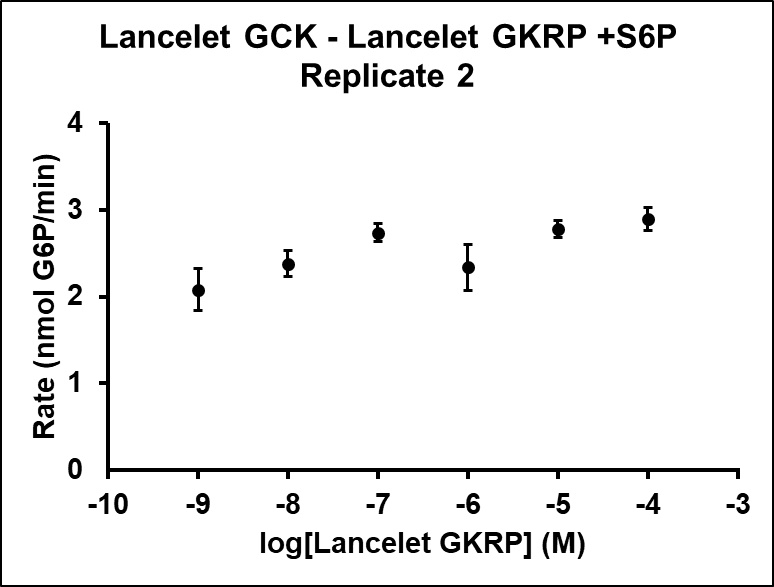

**Fig. S13:** Inhibition assay of Lancelet GCK with Lancelet GKRP. Each data point represents the average of triplicate measurements of the highest rate of glucose 6-phosphate production observed over at least one minute of data collection.

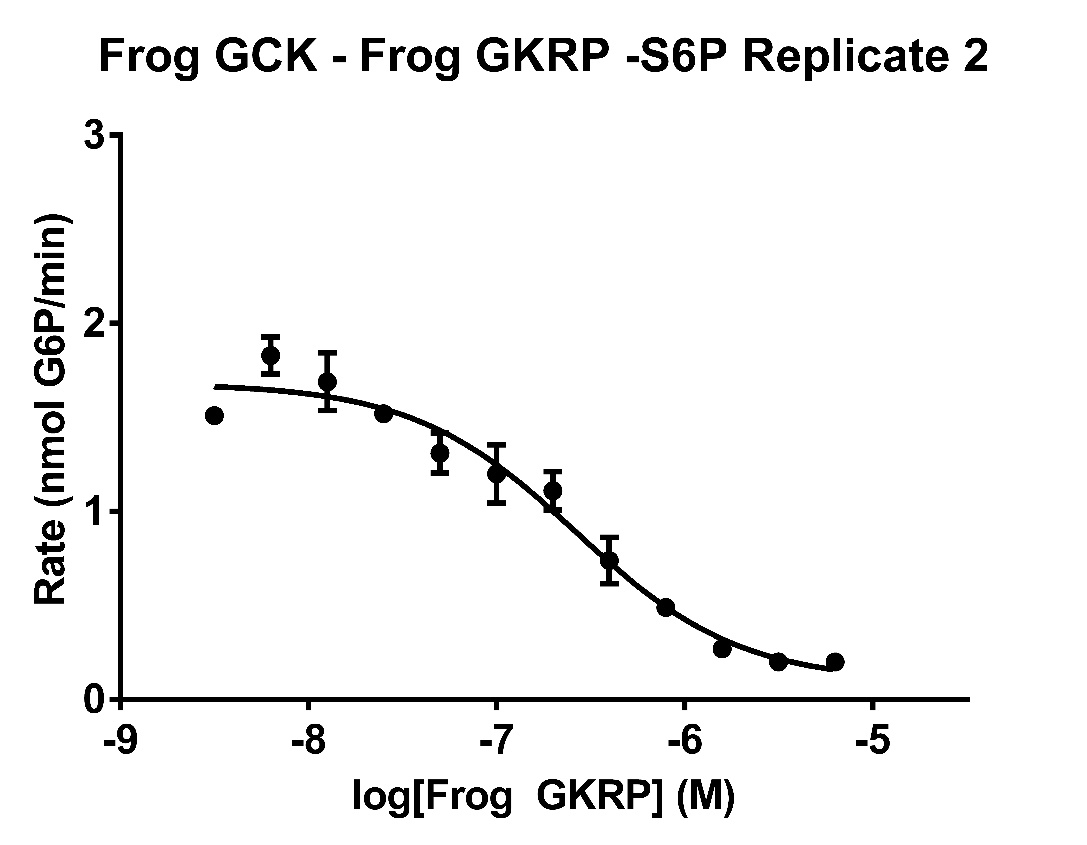

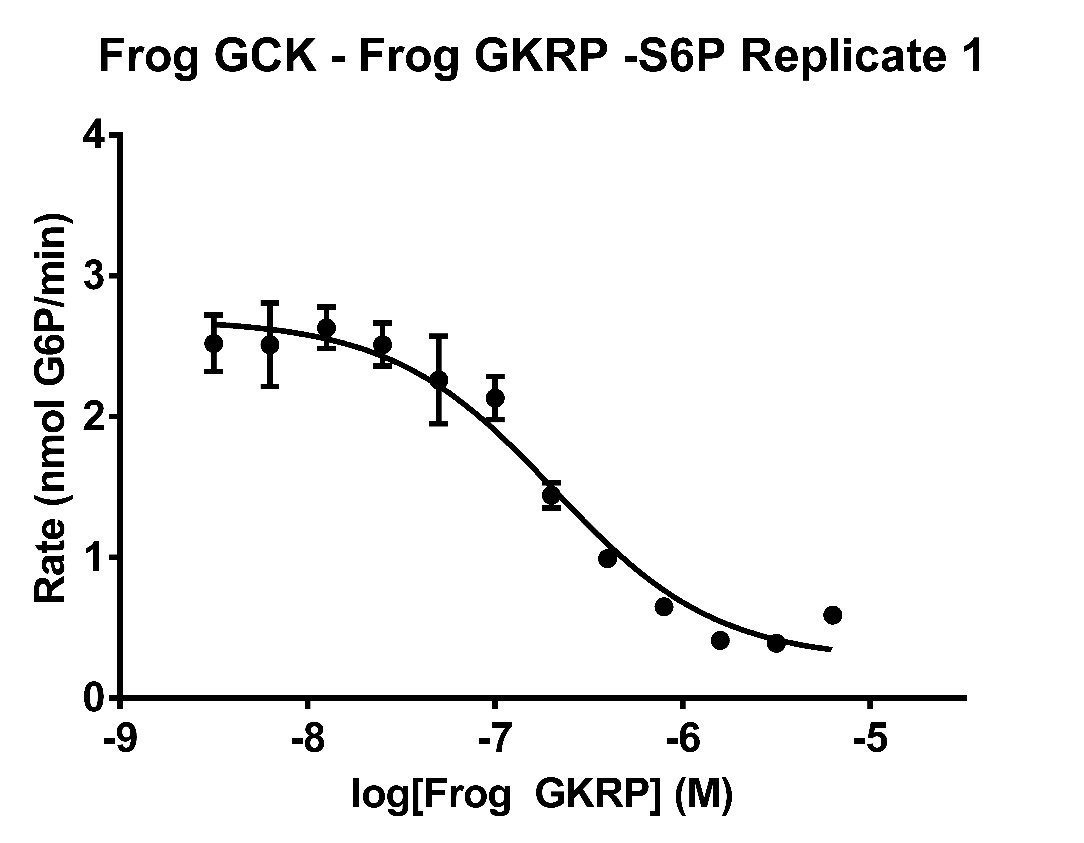

**Fig. S14:** Inhibition assay of Frog GCK with Frog GKRP in the absence of sugar phosphates. Each data point represents the average of triplicate measurements of the highest rate of glucose 6-phosphate production observed over at least one minute of data collection.

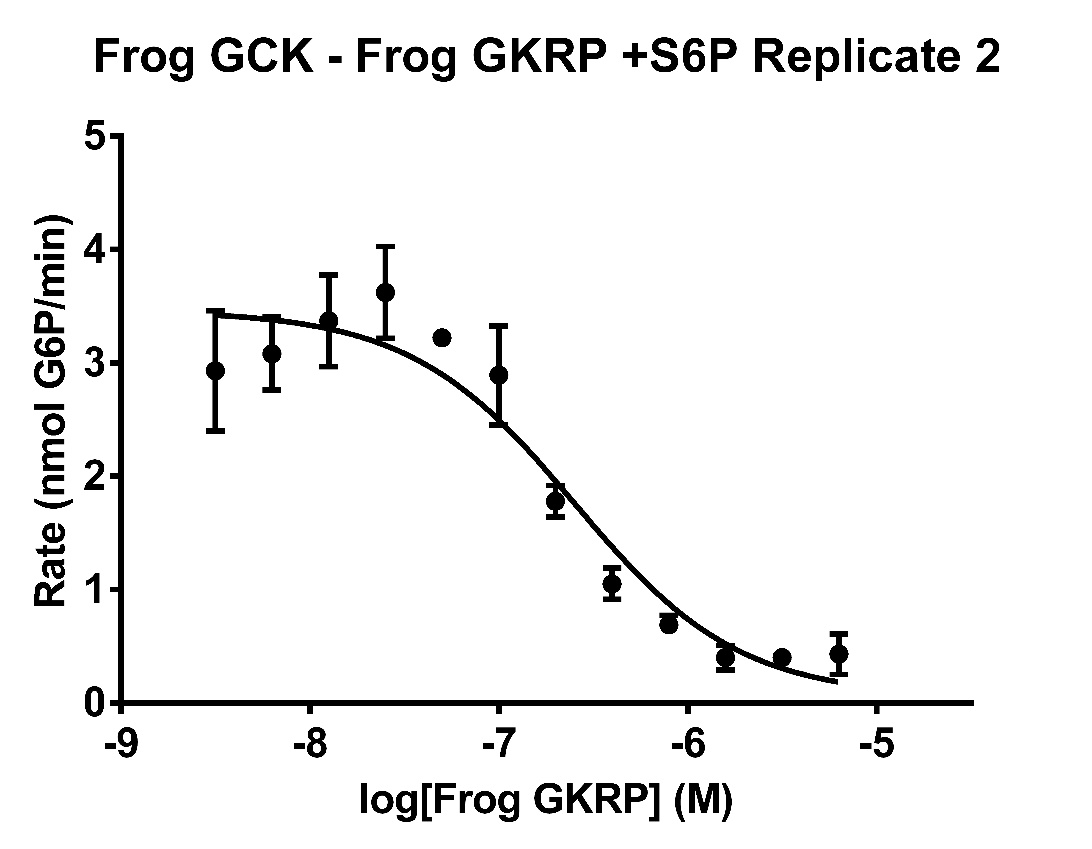

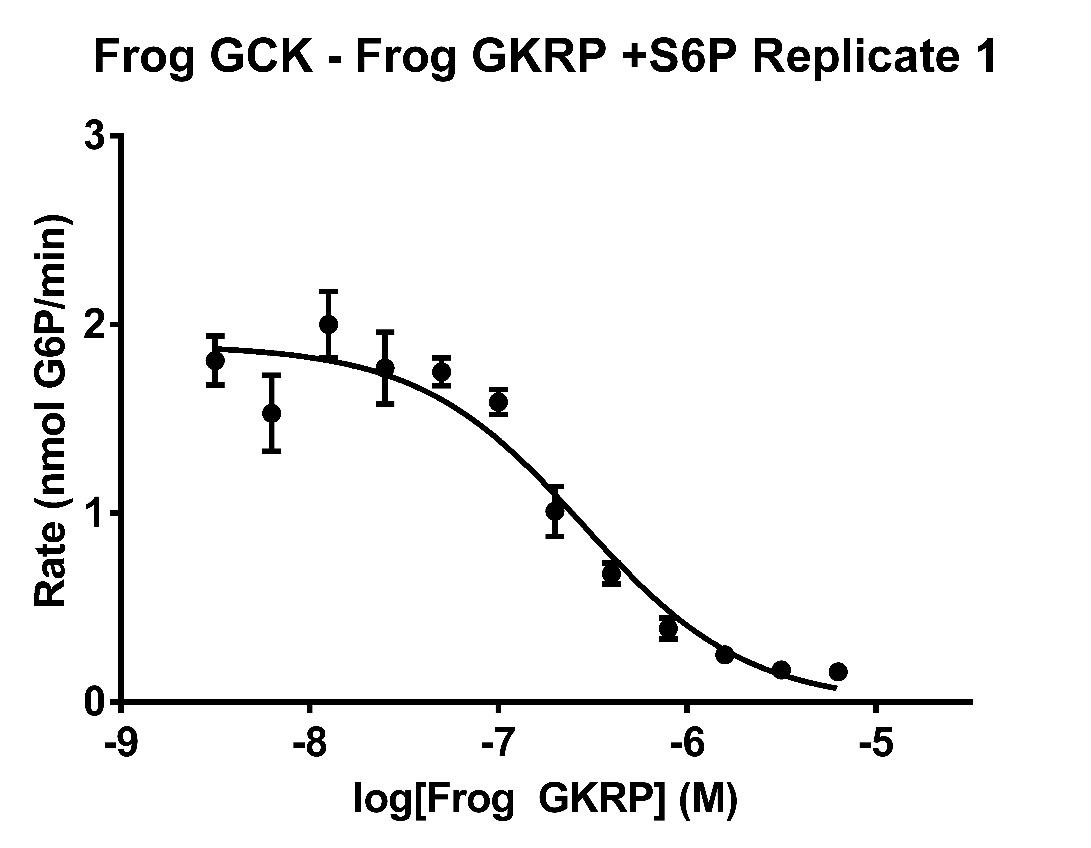

**Fig. S15:** Inhibition assay of Frog GCK with Frog GKRP in the presence of 2 mM S6P. Each data point represents the average of triplicate measurements of the highest rate of glucose 6-phosphate production observed over at least one minute of data collection.

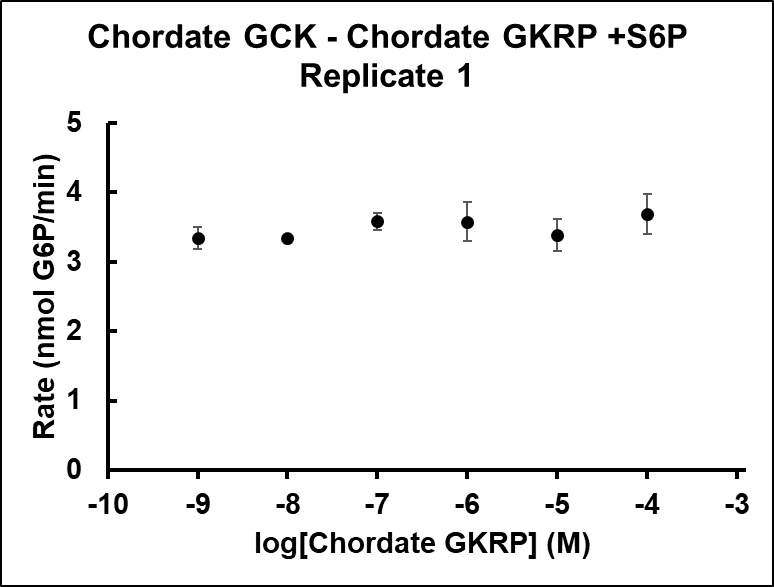

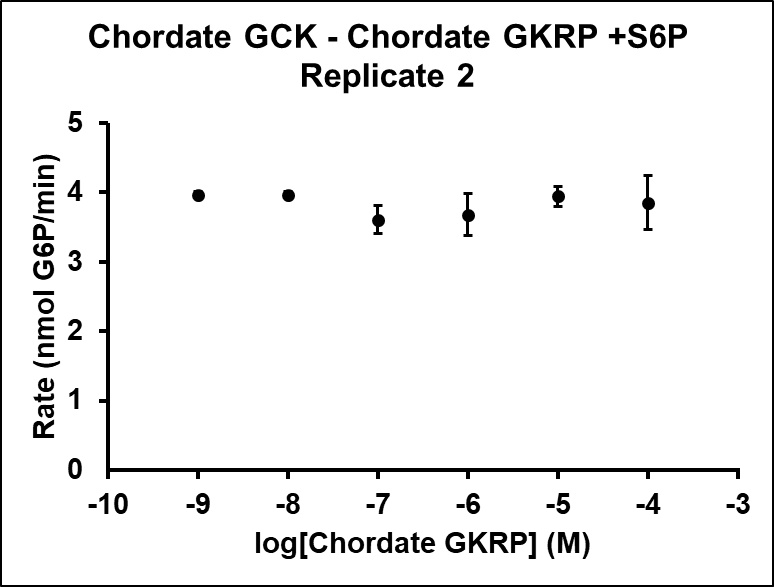

**Fig. S16:** Inhibition assay of Chordate GCK with Chordate GKRP. Each data point represents the average of triplicate measurements of the highest rate of glucose 6-phosphate production observed over at least one minute of data collection.

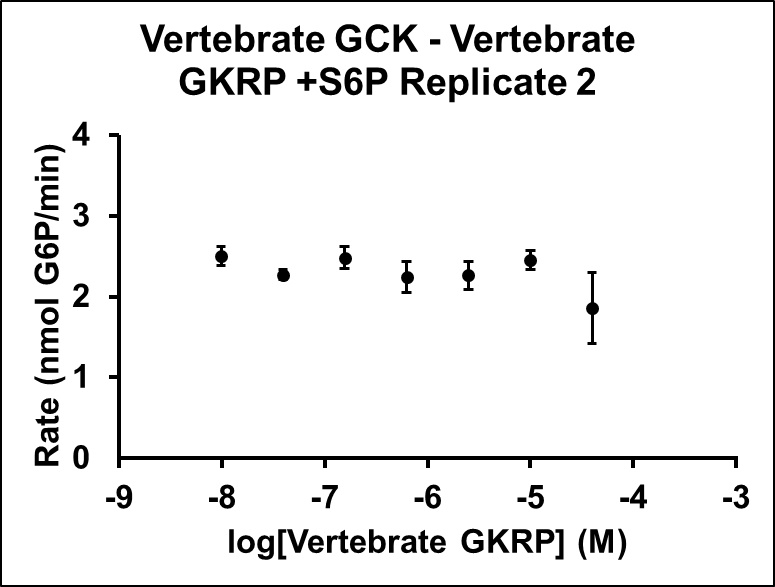

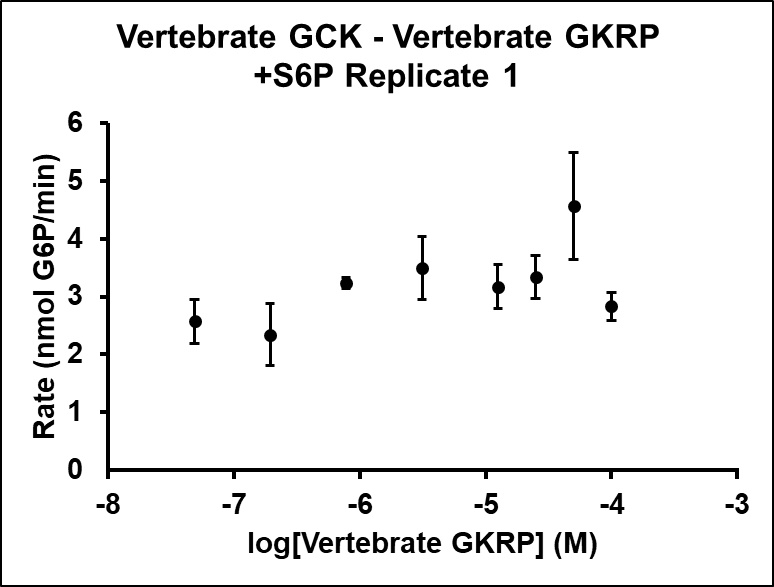

**Fig. S17:** Inhibition assay of Vertebrate GCK with Vertebrate GKRP. Each data point represents the average of triplicate measurements of the highest rate of glucose 6-phosphate production observed over at least one minute of data collection.

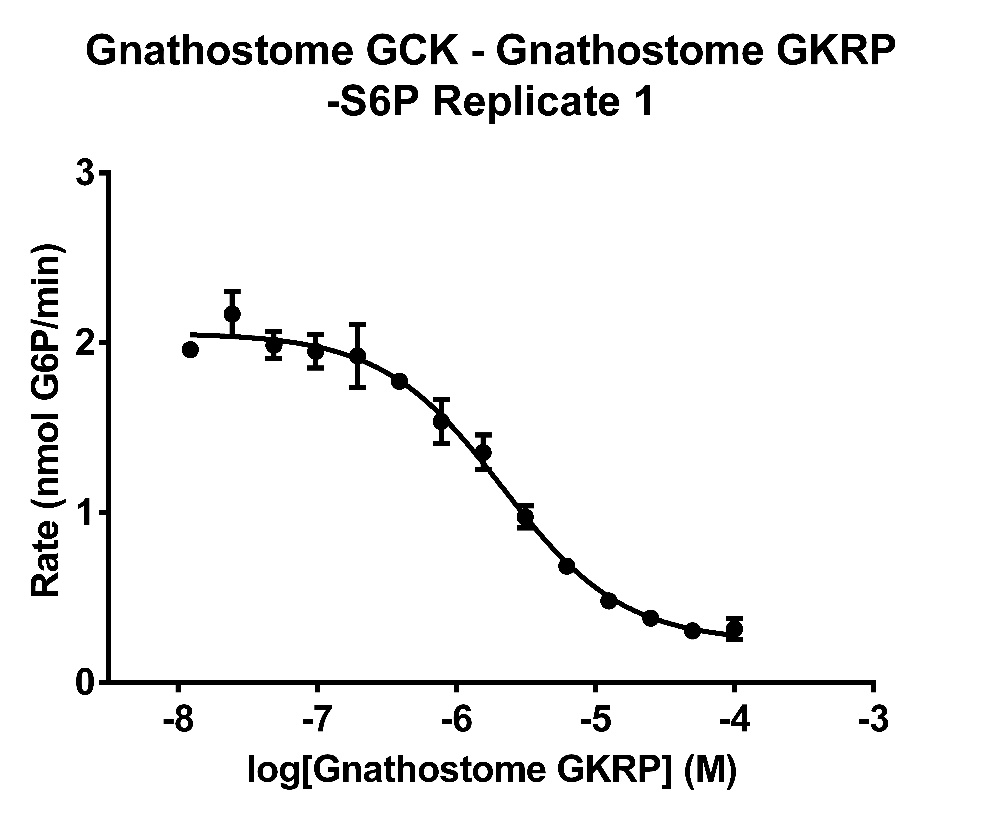

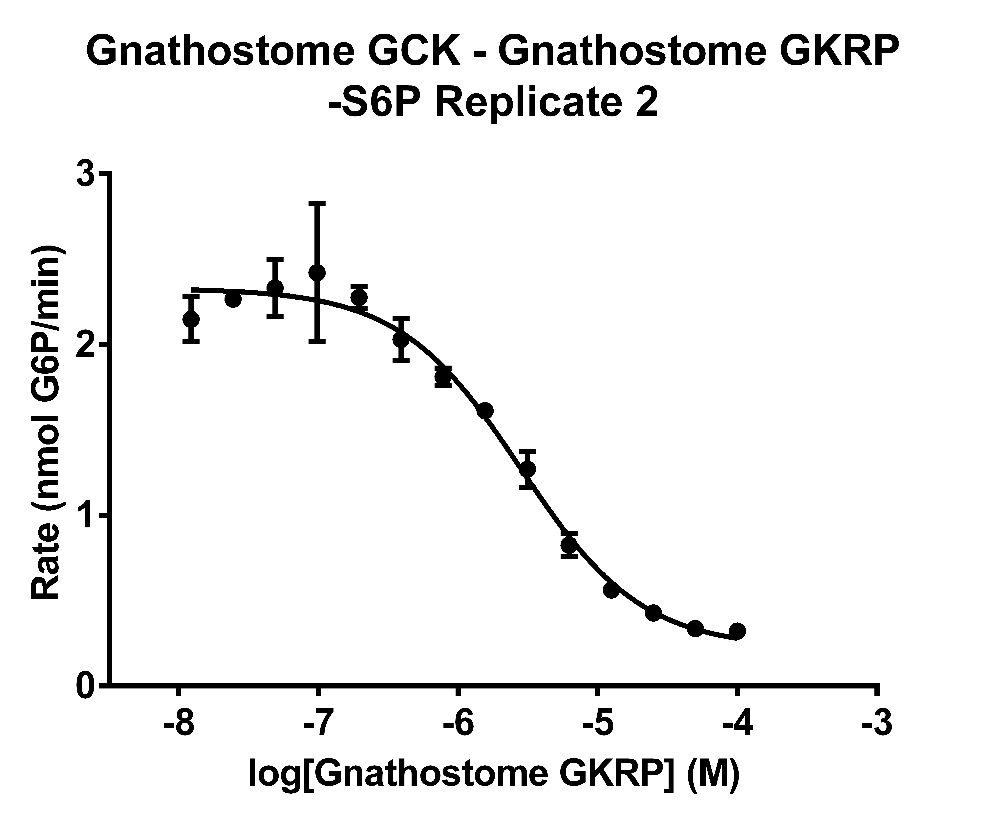

**Fig. S18:** Inhibition assay of Gnathostome GCK with Gnathostome GKRP in the absence of sugar phosphates. Each data point represents the average of triplicate measurements of the highest rate of glucose 6-phosphate production observed over at least one minute of data collection.

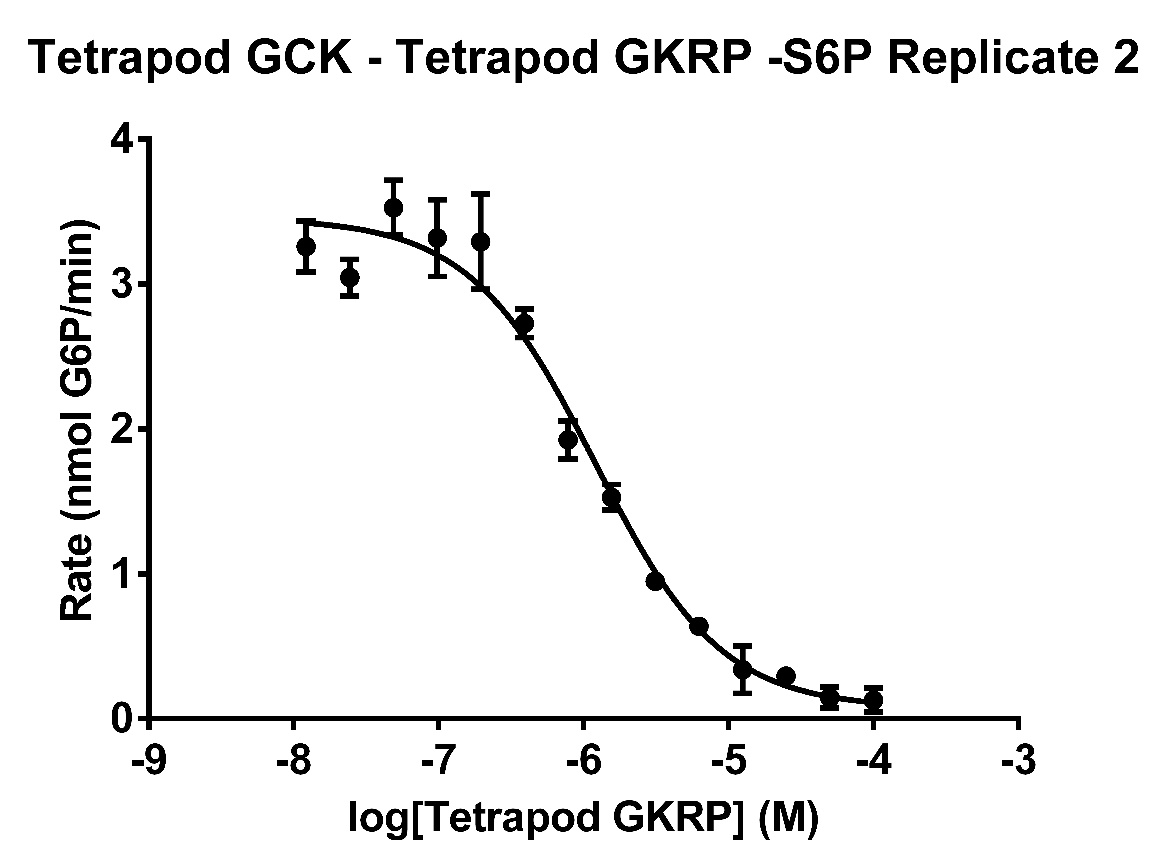

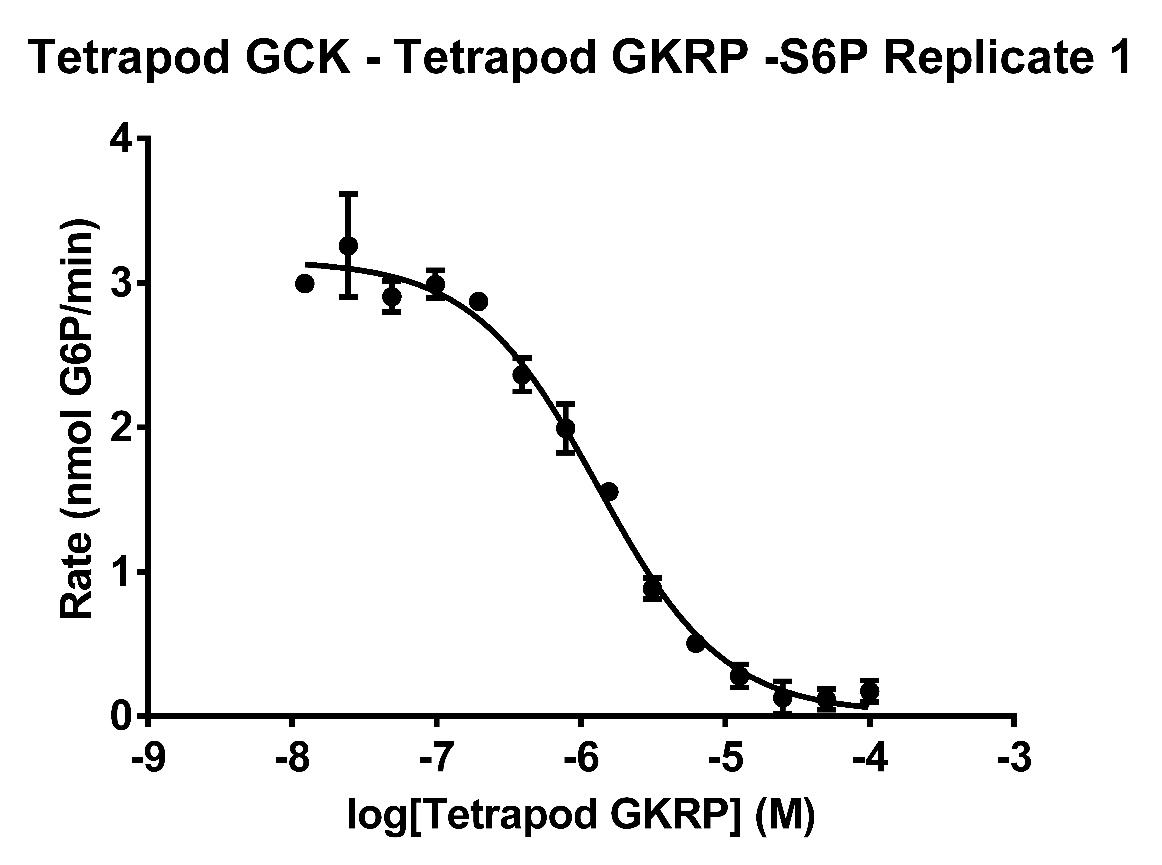

**Fig. S19:** Inhibition assay of Tetrapod GCK with Tetrapod GKRP in the absence of sugar phosphates. Each data point represents the average of triplicate measurements of the highest rate of glucose 6-phosphate production observed over at least one minute of data collection.

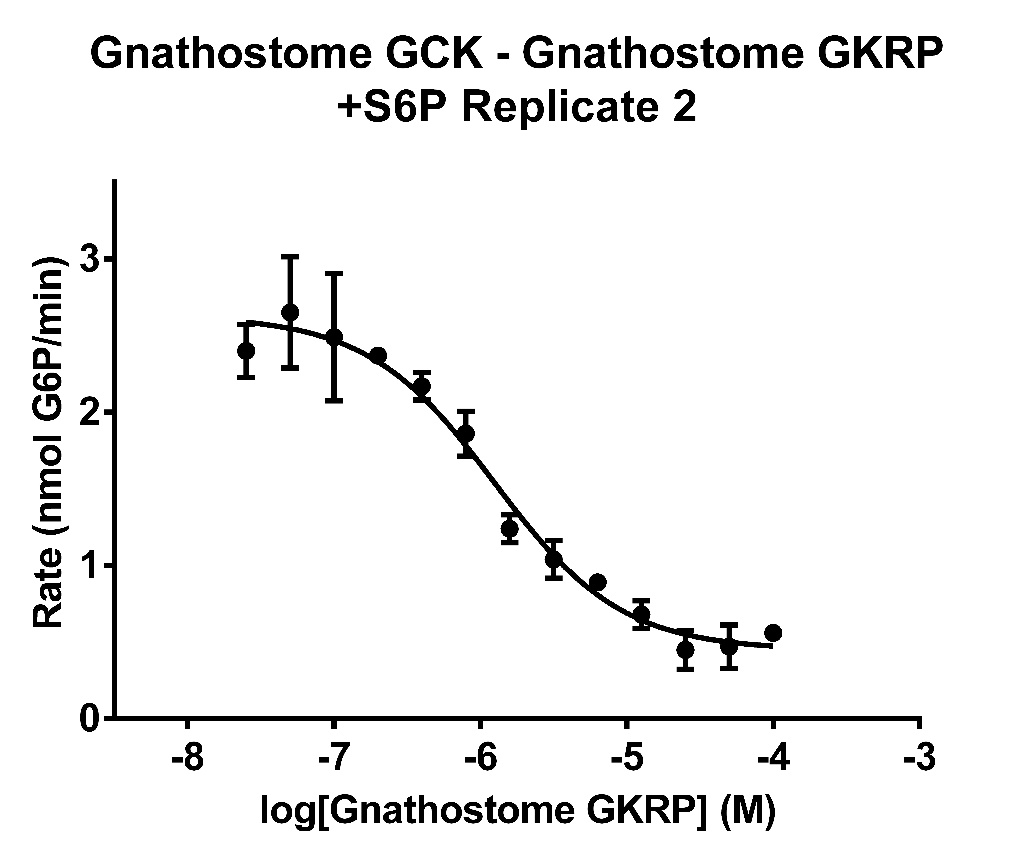

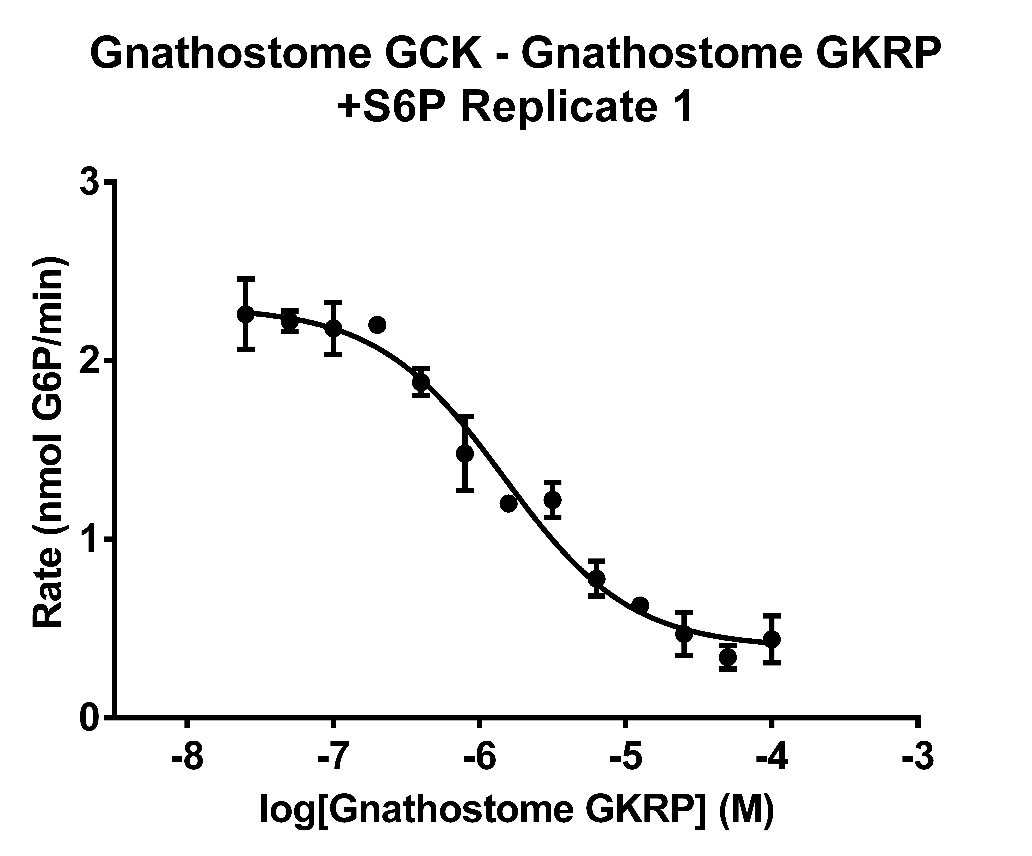

**Fig. S20:** Inhibition assay of Gnathostome GCK with Gnathostome GKRP in the presence of 2 mM S6P. Each data point represents the average of triplicate measurements of the highest rate of glucose 6-phosphate production observed over at least one minute of data collection.

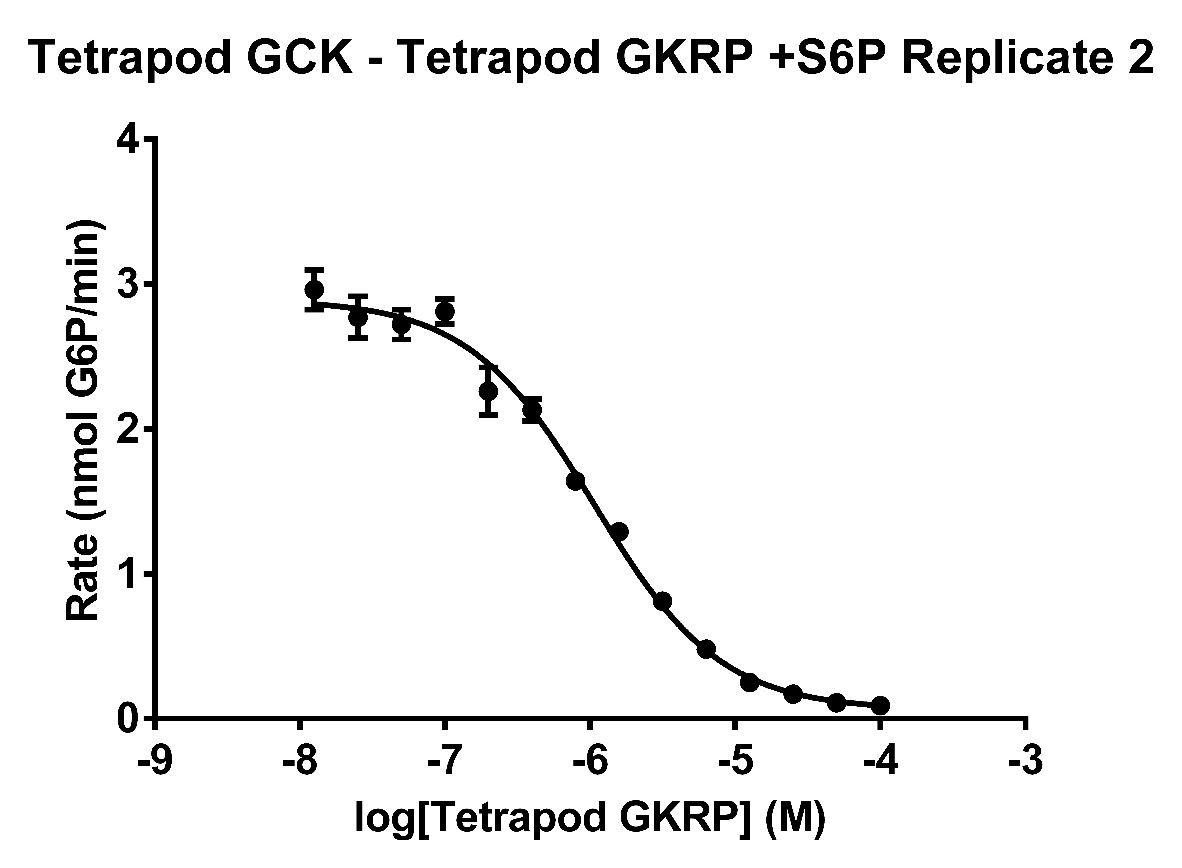

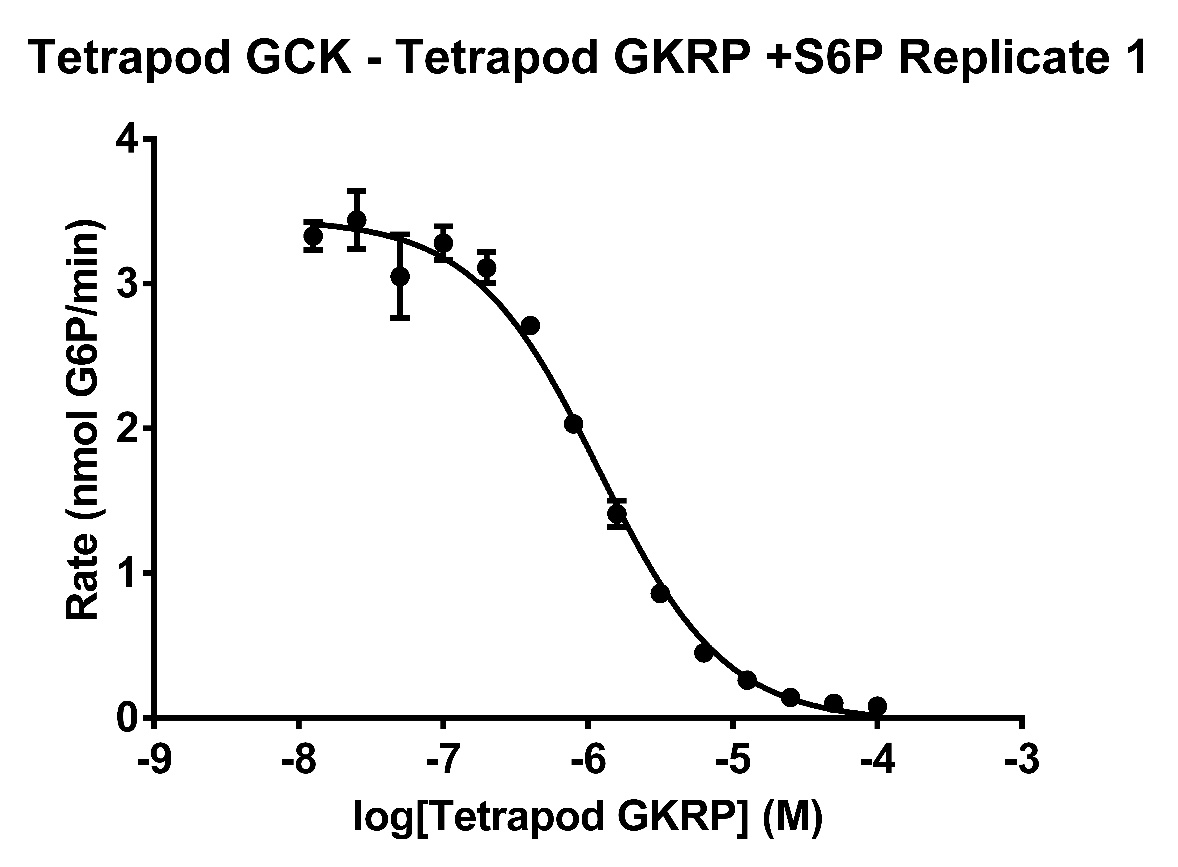

**Fig. S21:** Inhibition assay of Tetrapod GCK with Tetrapod GKRP in the presence of 2 mM S6P. Each data point represents the average of triplicate measurements of the highest rate of glucose 6-phosphate production observed over at least one minute of data collection.

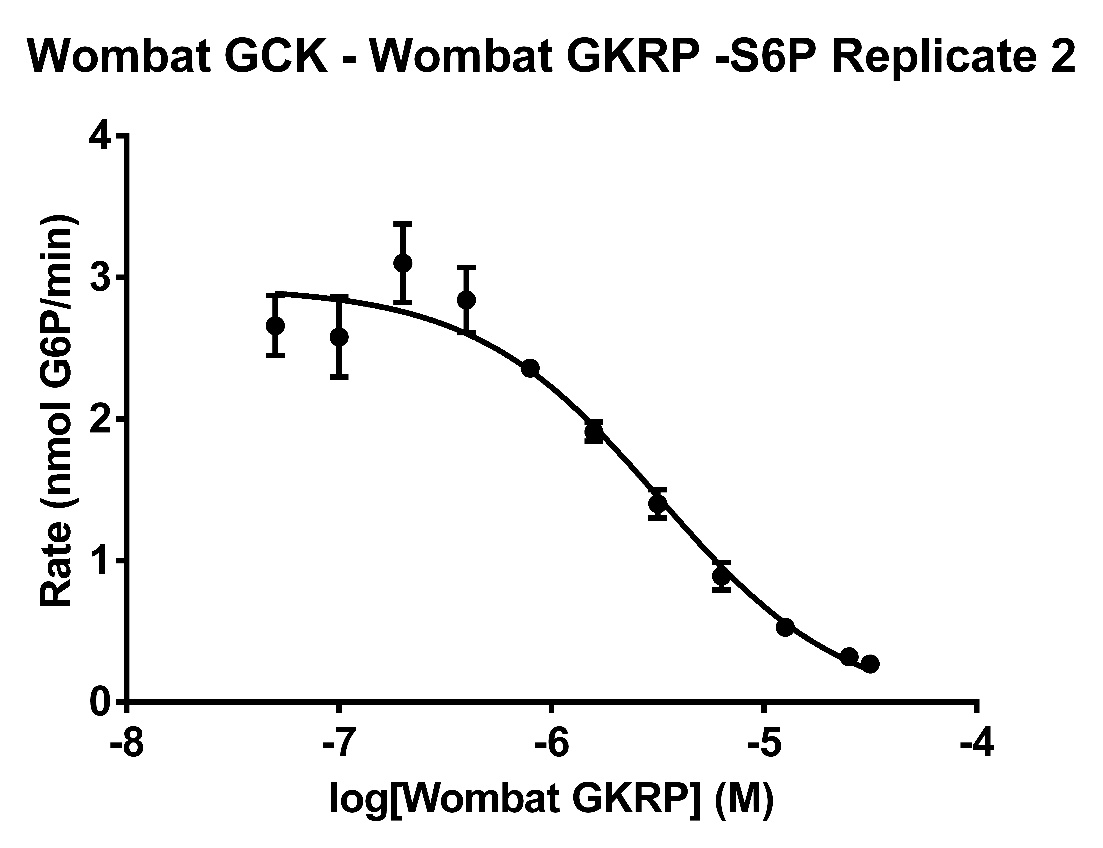

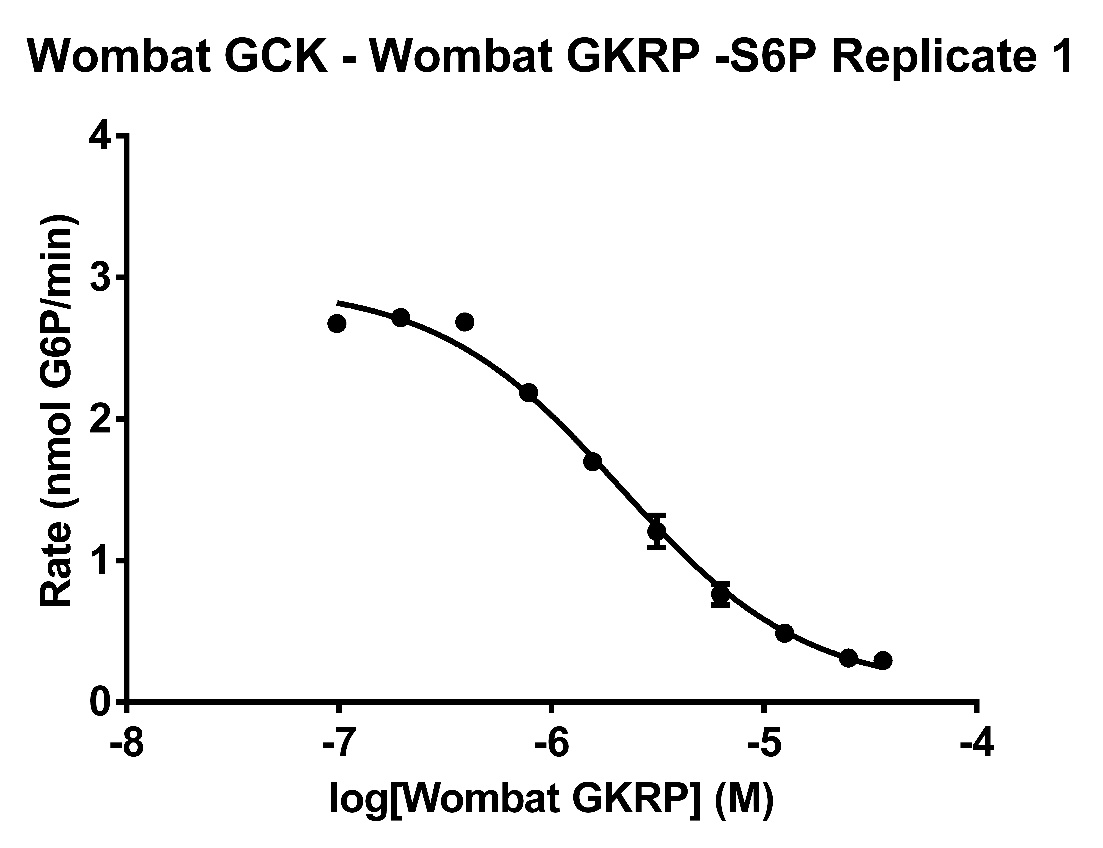

**Fig. S22:** Inhibition assay of Wombat GCK with Wombat GKRP in the absence of sugar phosphates. Each data point represents the average of triplicate measurements of the highest rate of glucose 6-phosphate production observed over at least one minute of data collection.

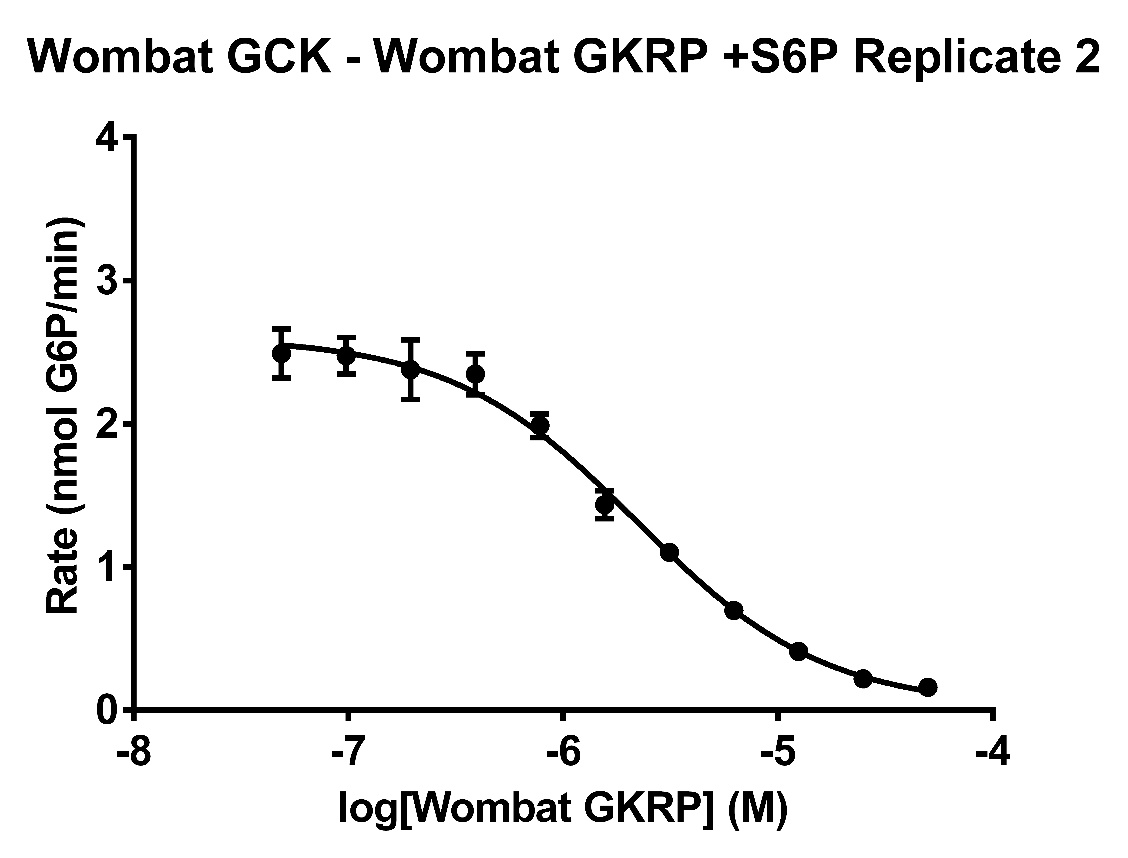

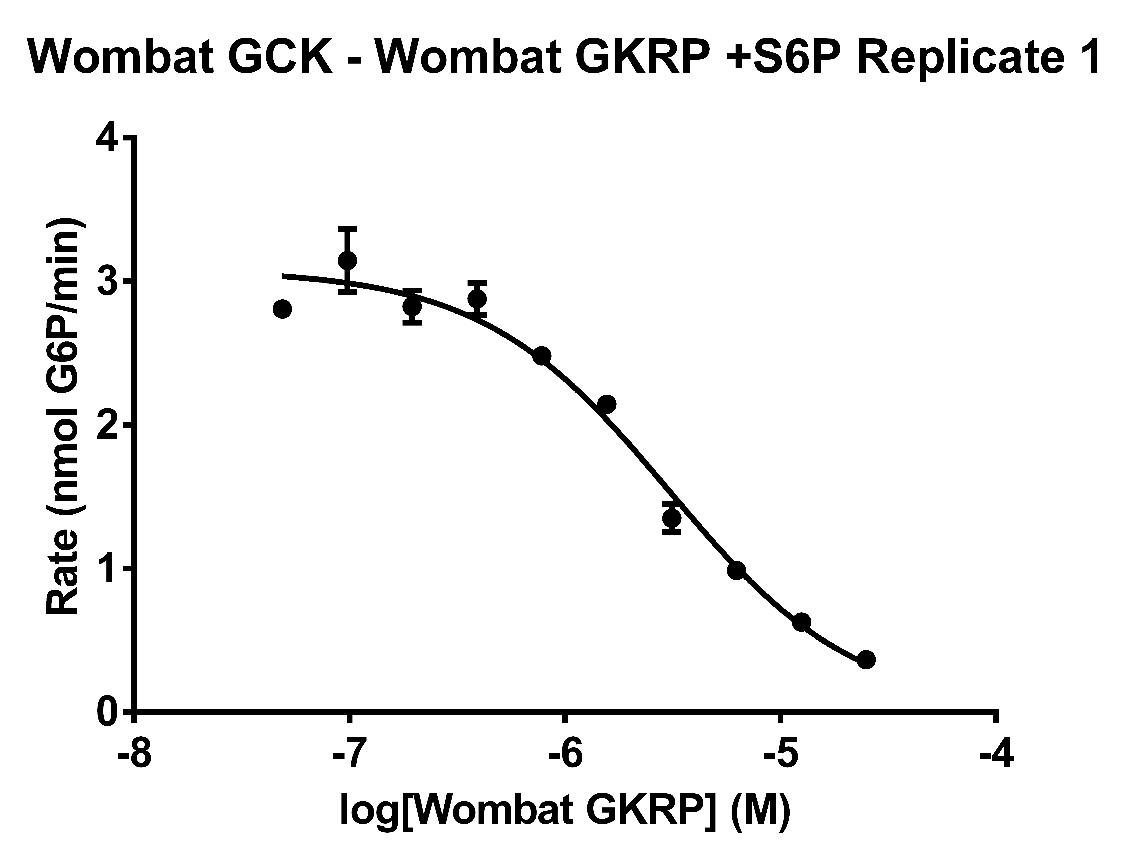

**Fig. S23:** Inhibition assay of Wombat GCK with Wombat GKRP in the presence of 2 mM S6P. Each data point represents the average of triplicate measurements of the highest rate of glucose 6-phosphate production observed over at least one minute of data collection.

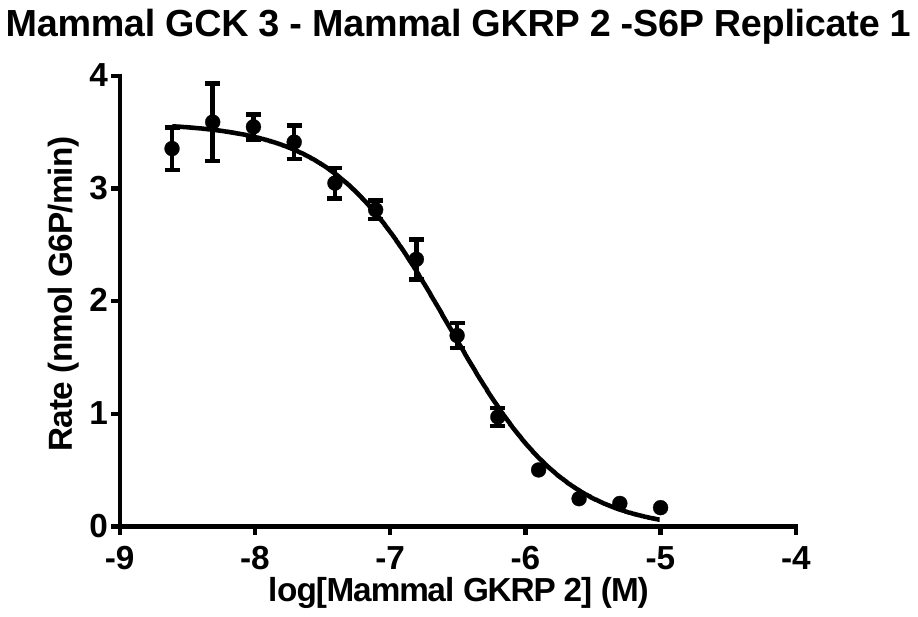

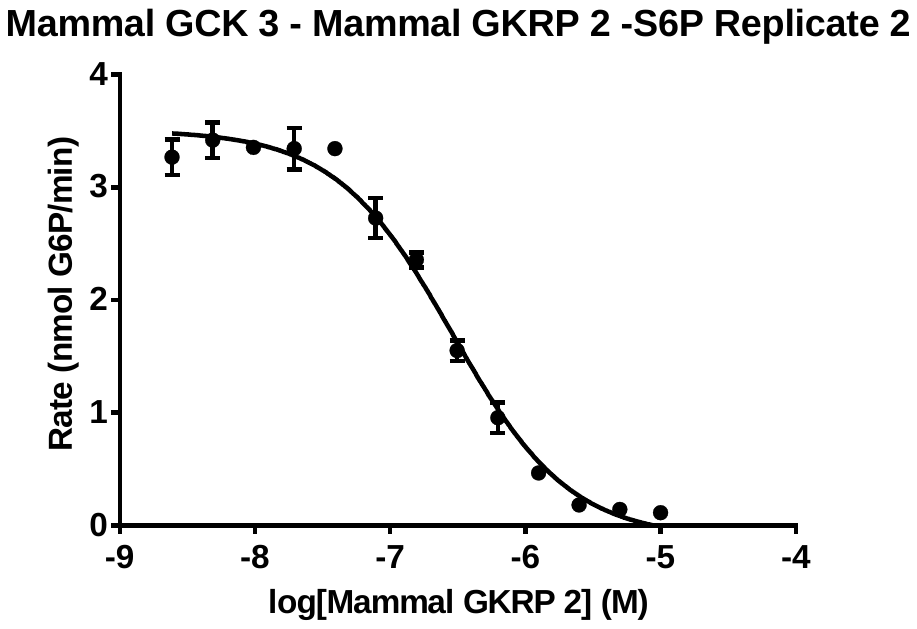

**Fig. S24:** Inhibition assay of M2 GCK with M1 GKRP in the absence of sugar phosphates. Each data point represents the average of triplicate measurements of the highest rate of glucose 6-phosphate production observed over at least one minute of data collection.

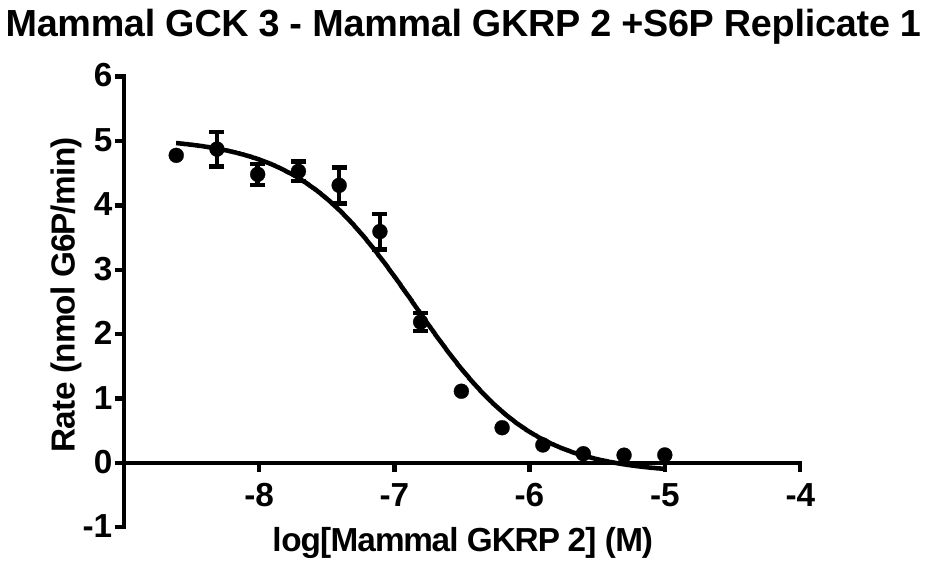

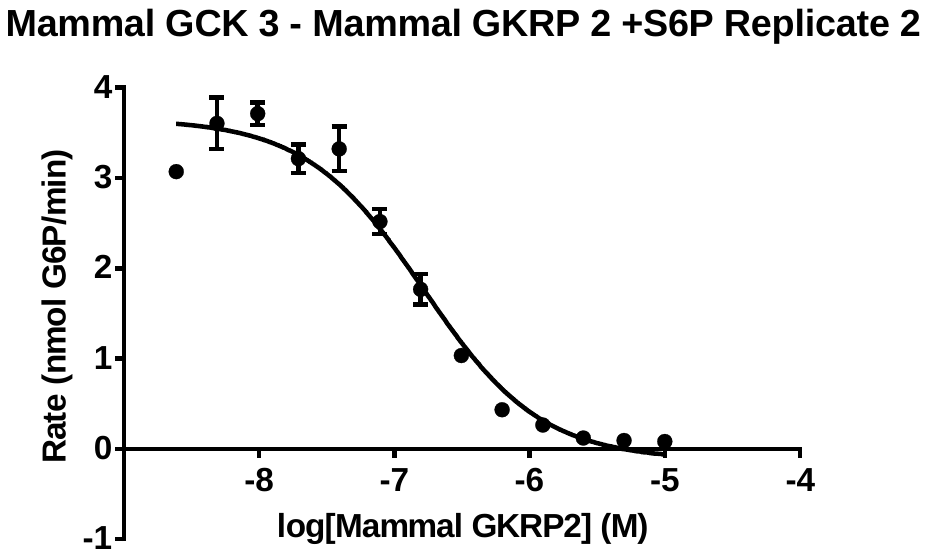

**Fig. S25:** Inhibition assay of M2 GCK with M1 GKRP in the presence of 2 mM S6P. Each data point represents the average of triplicate measurements of the highest rate of glucose 6-phosphate production observed over at least one minute of data collection.

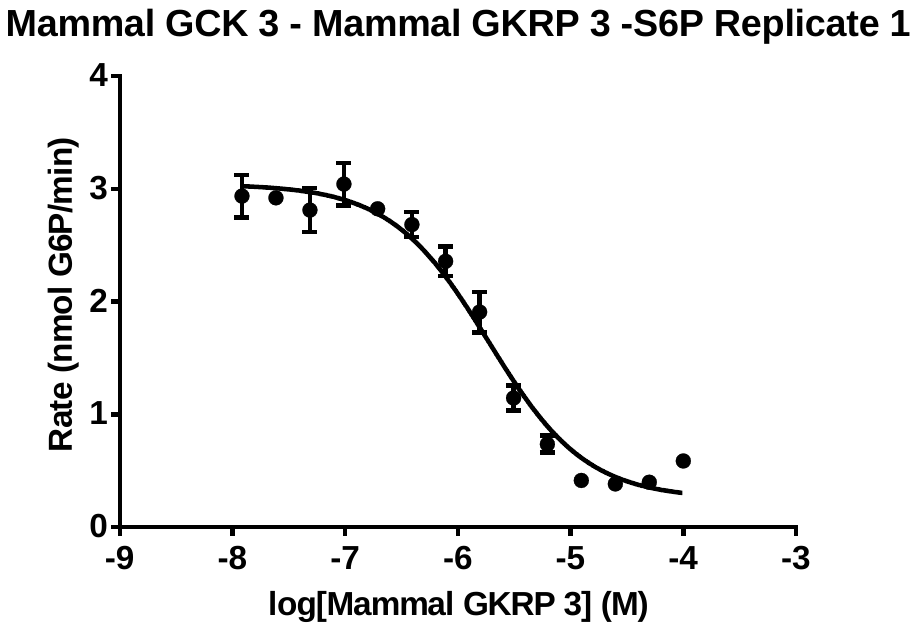

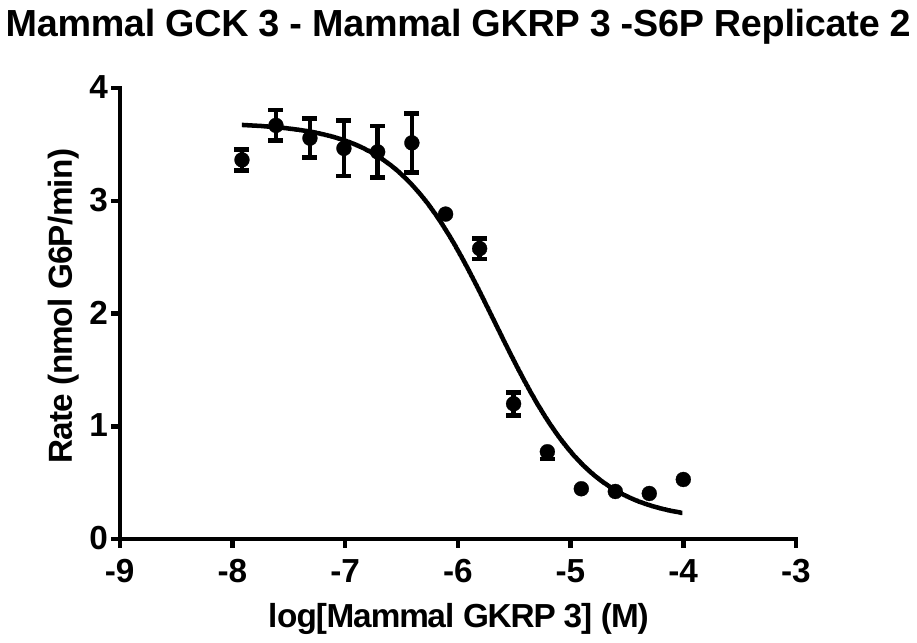

**Fig. S26:** Inhibition assay of M2 GCK with M2 GKRP in the absence of sugar phosphates. Each data point represents the average of triplicate measurements of the highest rate of glucose 6-phosphate production observed over at least one minute of data collection.

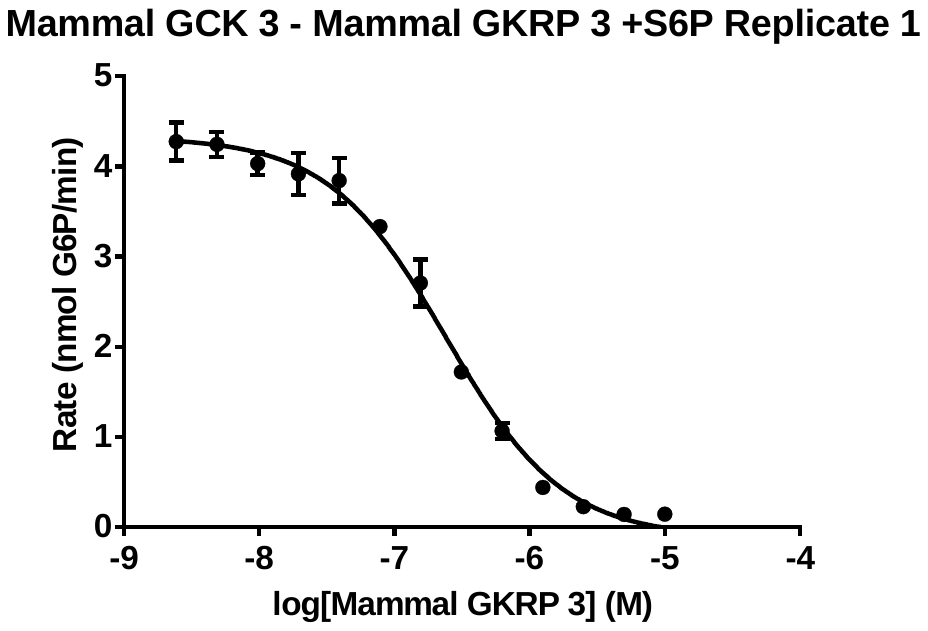

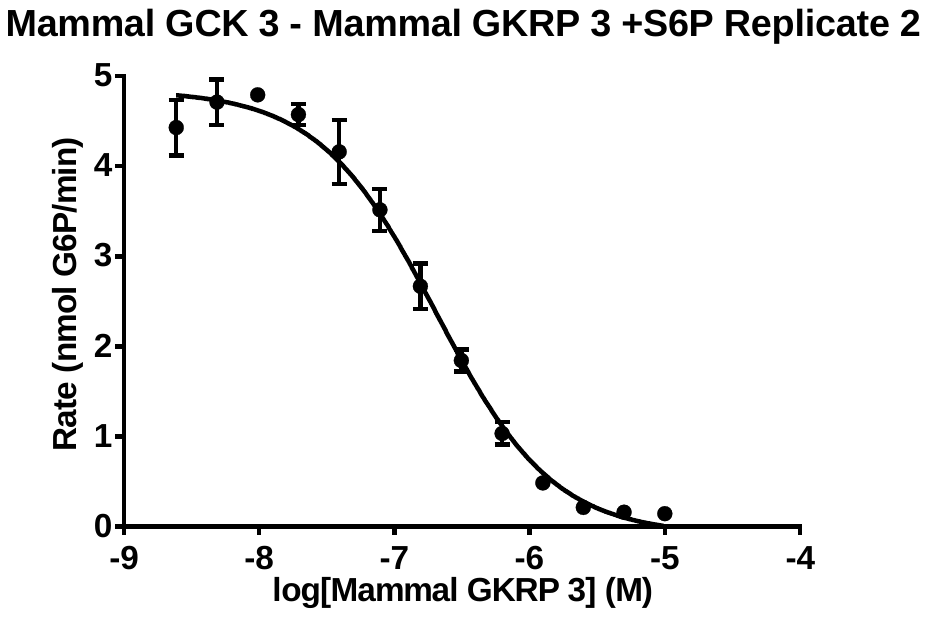

**Fig. S27:** Inhibition assay of M2 GCK with M2 GKRP in the presence of 2 mM S6P. Each data point represents the average of triplicate measurements of the highest rate of glucose 6-phosphate production observed over at least one minute of data collection.

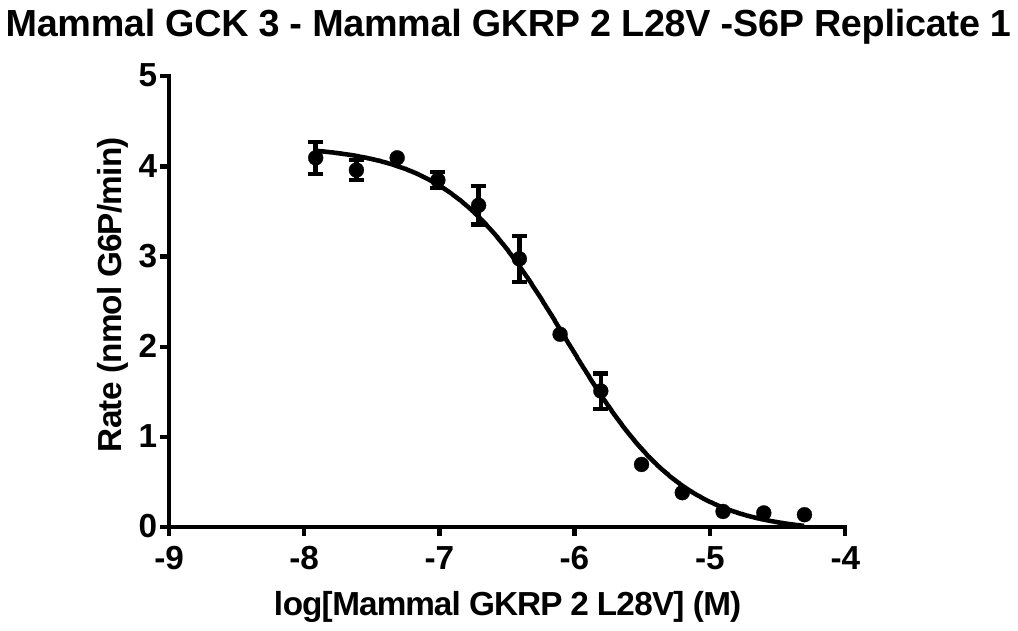

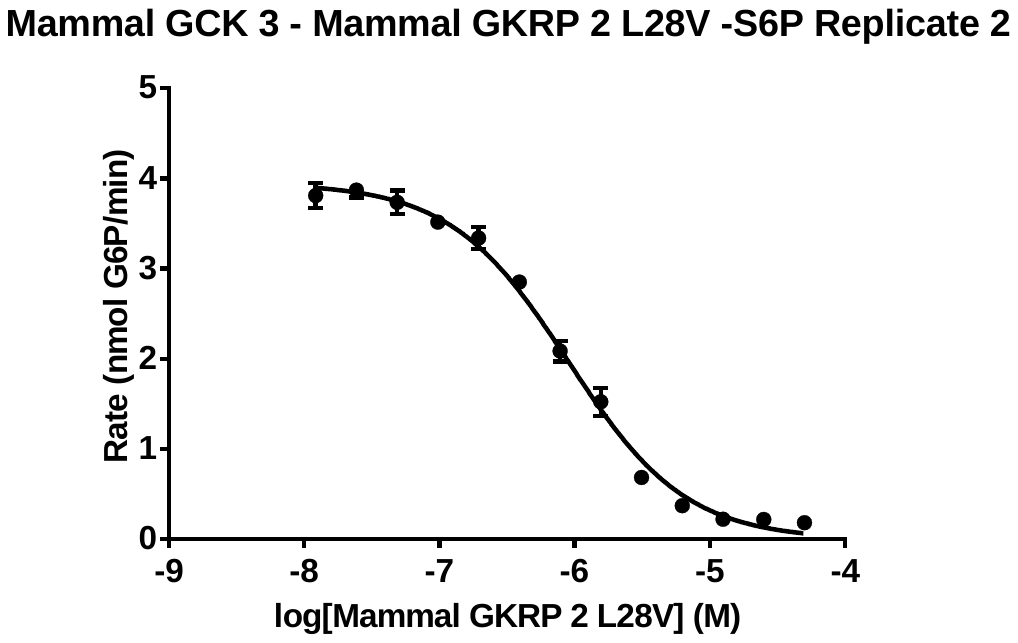

**Fig. S28:** Inhibition assay of Mammal GCK 3 with Mammal GKRP 2 L28V in the absence of sugar phosphates. Each data point represents the average of triplicate measurements of the highest rate of glucose 6-phosphate production observed over at least one minute of data collection.

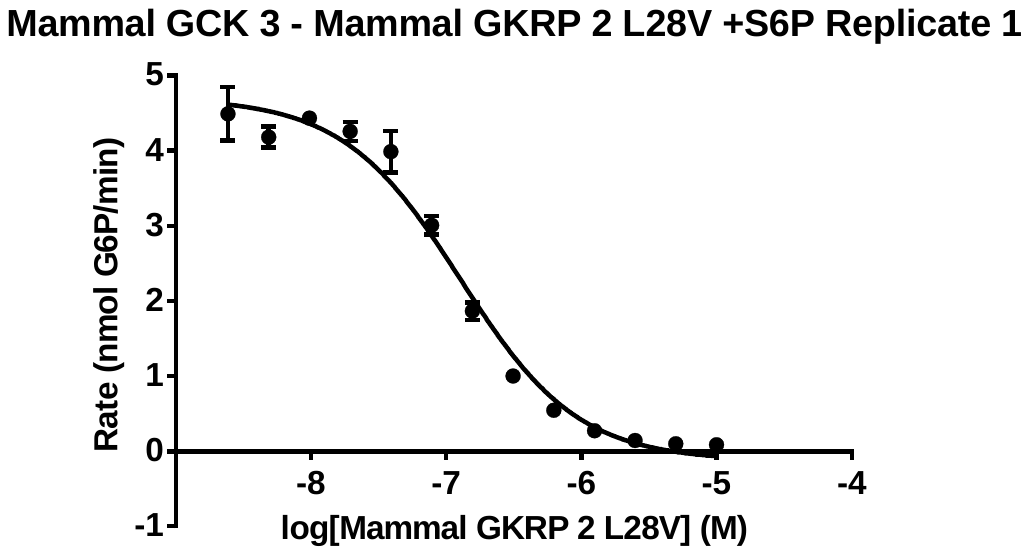

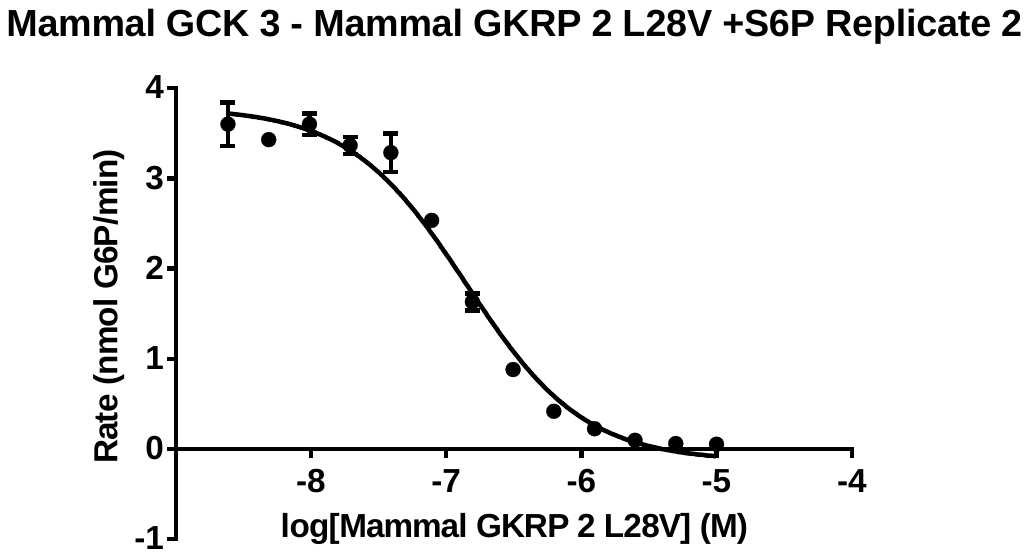

**Fig. S29:** Inhibition assay of Mammal GCK 3 with Mammal GKRP 2 L28V in the presence of 2 mM S6P. Each data point represents the average of triplicate measurements of the highest rate of glucose 6-phosphate production observed over at least one minute of data collection.

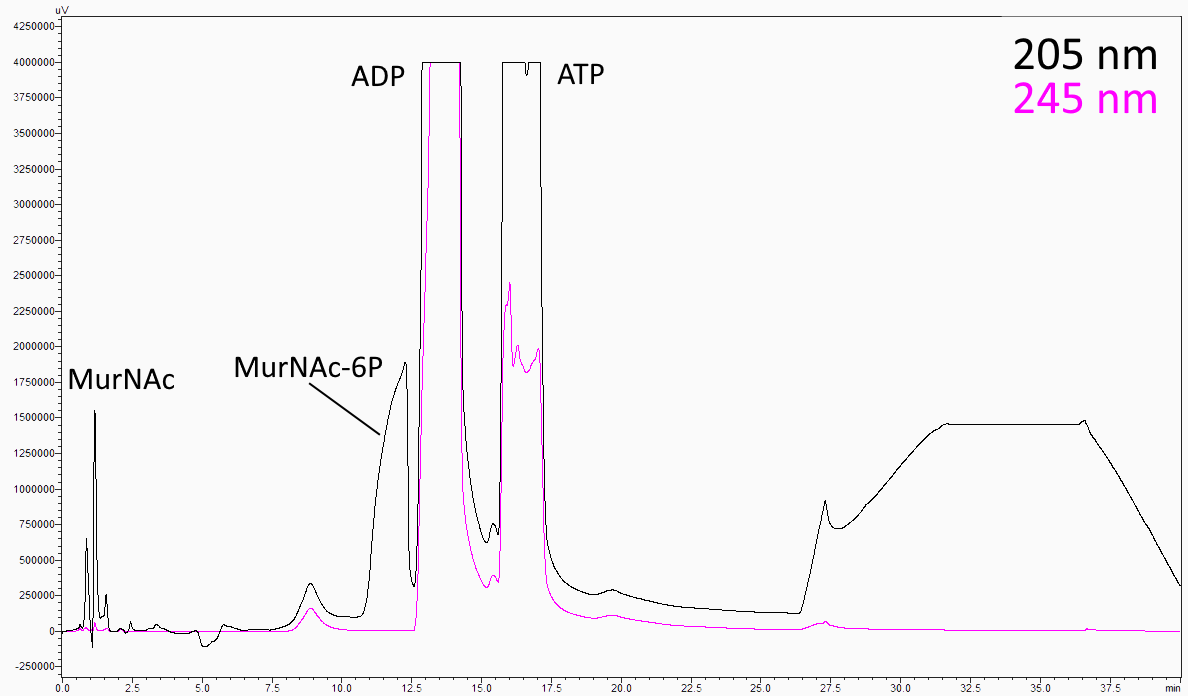

**Fig. S30:** HPLC chromatogram of the MurNAc-6P synthesis reaction. An example chromatogram produced during the separation of MurNAc-6P from the reaction mixture. The MurNAc, ADP, and ATP peaks were identified by loading each of those species separately.

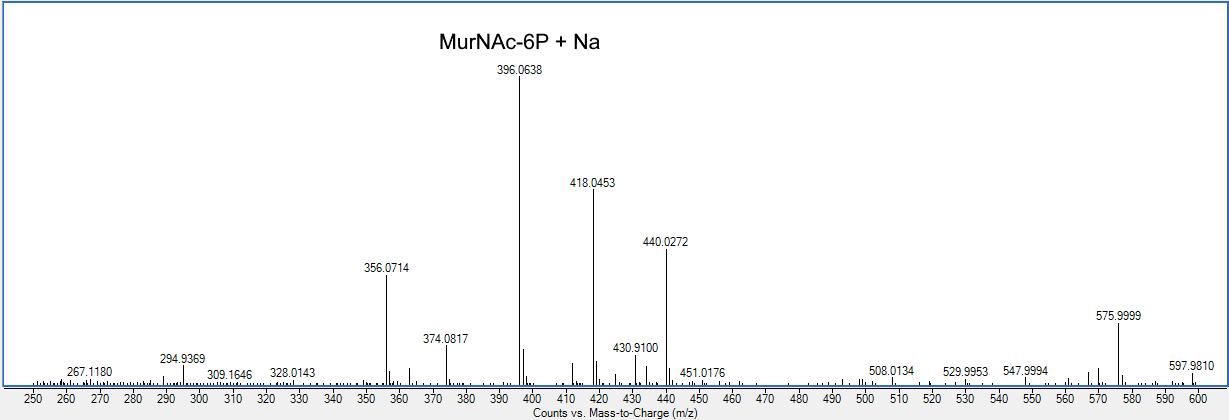

**Fig. S31:** Mass spectrum of purified MurNAc-6P. Peaks at 374, 396, 418, and 440 m/z represent [MurNAc-6P + H]^+^ with 0, 1, 2, or 3 Na^+^ ions bound, respectively. Fully protonated MurNAc-6P with a proton bound has a monoisotopic mass of 374.085227 Da.

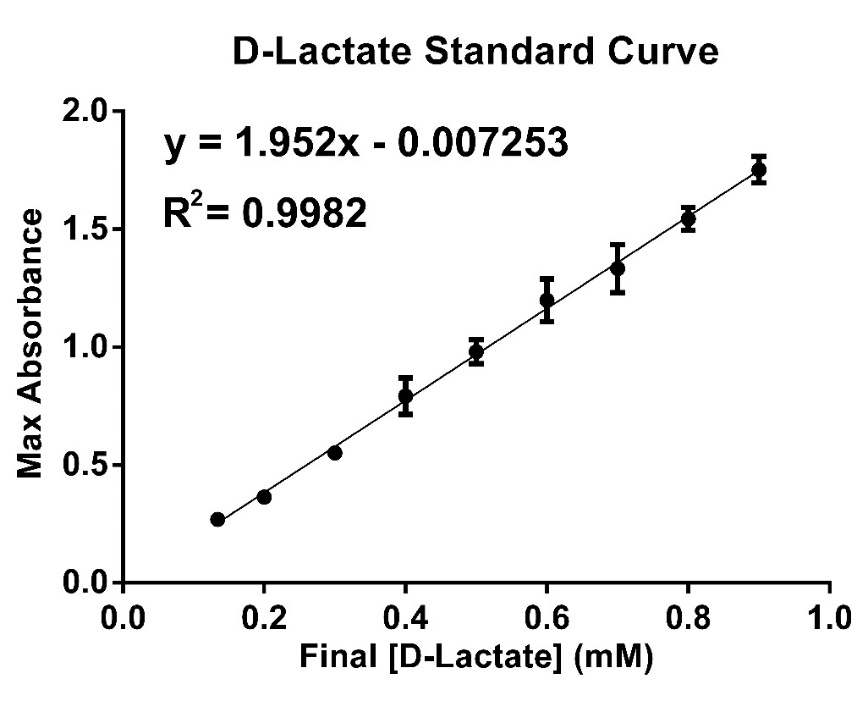

**Fig. S32**. D-lactate standard curve used to calculate the concentration of MurNAc-6P with the formazan indictor. Each data point represents the average of three technical replicates. Error bars represent standard deviation. Where not visible, the error bars are smaller than the data points.

**Table S1:** Kinetic parameters of the studied etherase activities.

| **Protein** | **Replicate** | ***k*_cat_ (s^-1^)** | ***K*_M_ (μM)** | ***k*_cat_/*K*_M_ (M^-1^ s^-1^)** | ***K*_I_ (mM)** |
| --- | --- | --- | --- | --- | --- |
| *E. coli* MurQ | 1 | 8.89 ± 1.45 | 338.4 ± 100.7 | 2.63 x 10^4^ ± 2.75 x 10^3^ | 1.91 ± 0.56 |
| *E. coli* MurQ | 2 | 1.11 ± 0.045 | 32.45 ± 5.06 | 3.14 x 10^4^ ± 7.35 x 10^3^ | 4.81 ± 0.72 |
| Rat GKRP | 1 | N/A | N/A | N/A | N/A |
| Rat GKRP | 2 | N/A | N/A | N/A | N/A |
| Lancelet GKRP | 1 | 0.367 ± 0.0055 | 198.1 ± 13.2 | 1853.6 ± 89.8 | N/A |
| Lancelet GKRP | 2 | 0.441 ± 0.016 | 389.8 ± 45.8 | 1131.6 ± 196.3 | N/A |
| Chordate GKRP | 1 | 0.211 ± 0.0083 | 276.8 ± 38.8 | 762 ± 67.1 | N/A |
| Chordate GKRP | 2 | 0.156 ± 0.007 | 242.1 ± 38.6 | 644.4 ± 63.7 | N/A |
| Vertebrate GKRP | 1 | 0.0190 ± 0.0011 | 215.9 ± 46.4 | 88.0 ± 1.48 | N/A |
| Vertebrate GKRP | 2 | 0.0183 ± 0.0004 | 219.1 ± 18.6 | 83.5 ± 9.7 | N/A |
| Gnathostome GKRP | 1 | 0.0210 ± 0.0006 | 250.2 ± 33.3 | 83.9 ± 7.7 | N/A |
| Gnathostome GKRP | 2 | 0.0262 ± 0.0008 | 207.7 ± 23.8 | 126.1 ± 20.7 | N/A |

**Table S2:** IC_50_ values of the studied GCK-GKRP interactions.

| Protein | (+/-) S6P | Replicate | IC_50_ (μM) |
| --- | --- | --- | --- |
| Lancelet GKRP | + | 1 | N/A |
| Lancelet GKRP | + | 2 | N/A |
| Frog GKRP | - | 1 | 0.201 ± 0.054 |
| Frog GKRP | - | 2 | 0.263 ± 0.0728 |
| Frog GKRP | + | 1 | 0.247 ± 0.111 |
| Frog GKRP | + | 2 | 0.273 ± 0.101 |
| Wombat GKRP | - | 1 | 2.10 ± 0.33 |
| Wombat GKRP | - | 2 | 3.21 ± 1.03 |
| Wombat GKRP | + | 1 | 3.03 ± 0.74 |
| Wombat GKRP | + | 2 | 2.22 ± 0.24 |
| Chordate GKRP | + | 1 | N/A |
| Chordate GKRP | + | 2 | N/A |
| Vertebrate GKRP | + | 1 | N/A |
| Vertebrate GKRP | + | 2 | N/A |
| Gnathostome GKRP | - | 1 | 2.15 ± 0.21 |
| Gnathostome GKRP | - | 2 | 2.8 ± 0.04 |
| Gnathostome GKRP | + | 1 | 1.45 ± 0.25 |
| Gnathostome GKRP | + | 2 | 2.12 ± 0.55 |
| Tetrapod GKRP | - | 1 | 1.32 ± 0.13 |
| Tetrapod GKRP | - | 2 | 1.21 ± 0.19 |
| Tetrapod GKRP | + | 1 | 1.05 ± 0.10 |
| Tetrapod GKRP | + | 2 | 1.17 ± 0.12 |
| Mammal GKRP 2 | - | 1 | 0.274 ± 0.0297 |
| Mammal GKRP 2 | - | 2 | 0.29 ± 0.04 |
| Mammal GKRP 2 | + | 1 | 0.142 ± 0.0262 |
| Mammal GKRP 2 | + | 2 | 0.16 ± 0.04 |
| Mammal GKRP 2 L28V | - | 1 | 0.86 ± 0.08 |
| Mammal GKRP 2 L28V | - | 2 | 0.90 ± 0.09 |
| Mammal GKRP 2 L28V | + | 1 | 0.13 ± 0.02 |
| Mammal GKRP 2 L28V | + | 2 | 0.14 ± 0.02 |
| Mammal GKRP 3 | - | 1 | 1.87 ± 0.39 |
| Mammal GKRP 3 | - | 2 | 2.13 ± 0.597 |
| Mammal GKRP 3 | + | 1 | 0.24 ± 0.0222 |
| Mammal GKRP 3 | + | 2 | 0.203 ± 0.0232 |

1.	S. S. Kamalaldinezabadi, *et al.*, Evolution of Protein Regulation in the Vertebrate Glucose Sensor. *bioRxiv* 2026.05.05.723016 (2026). https://doi.org/10.64898/2026.05.05.723016.

2.	S. Unsleber, M. Borisova, C. Mayer, Enzymatic synthesis and semi-preparative isolation of N-acetylmuramic acid 6-phosphate. *Carbohydr. Res.* **445**, 98–103 (2017).
